# Dissecting the roles of Expansion/Rebuf and the chitin synthase Krotzkopf Verkehrt in chitin deposition in *Drosophila*

**DOI:** 10.1101/2022.07.21.500966

**Authors:** Ettore De Giorgio, Panagiotis Giannios, M. Lluisa Espinàs, Marta Llimargas

## Abstract

Chitin is a highly abundant polymer in nature and a principal component of apical extracellular matrices in insects. In addition, chitin has proved to be an excellent biomaterial with multiple applications. In spite of its importance, the molecular mechanisms of chitin biosynthesis and chitin structural diversity are not fully elucidated yet. To investigate these issues, we use *Drosophila* as a model. We previously showed that chitin deposition in ectodermal tissues requires the concomitant activities of the chitin synthase enzyme Kkv and the functionally interchangeable proteins Exp and Reb. Here we carry out a cellular and molecular analysis of chitin deposition and we show that chitin polymerisation and chitin translocation to the extracellular space are uncoupled. We find that Kkv activity in chitin translocation, but not in polymerisation, requires the activity of Exp/Reb, and in particular of its conserved Nα-MH2 domain. The activity of Kkv in chitin polymerisation and translocation correlate with Kkv subcellular localisation, and in absence of Kkv-mediated extracellular chitin deposition, chitin accumulates intracellularly as membrane-less punctae. Furthermore, we find that Kkv and Exp/Reb display a largely complementary pattern at the apical domain, and that Exp/Reb activity regulates the topological distribution of Kkv at the apical membrane. We propose a model in which Exp/Reb regulates the organisation of Kkv complexes at the apical membrane which, in turn, regulates the function of Kkv in extracellular chitin translocation.

## INTRODUCTION

Chitin, a polymer of UDP-N-acetylglucosamine (GlcNAc) monomers, is a principal component of the apical extracellular matrix in arthropods. Chitin has a recognised importance in physiology (Zhao, 2019; Zhu et al., 2016) but also as a biomaterial. Chitin and its deacetylated form, chitosan, are non-toxic and biodegradable biopolymers with numerous applications in many sectors such as biomedicine, biotechnology, water treatment, food, agriculture, veterinary, or cosmetics (Casadidio et al., 2019; Elieh-Ali-Komi and Hamblin, 2016). So far, the main commercial sources of chitin are crab and shrimp shells (Younes and Rinaudo, 2015). Chitin isolation and purification from these sources require several treatments to remove proteins, calcium carbonate, lipids and pigments, and no standarised methods exists nowadays (Younes and Rinaudo, 2015). In addition, these treatments have many industrial drawbacks such as high energy consumption, long handling times, solvent waste, high environmental pollution and high economical costs, among others (Casadidio et al., 2019). The synthesis of chitin *in vitro* can represent a more ecological, efficient and “green” method as an alternative to the chemical procedures. Thus, it is critical to understand the molecular mechanisms of chitin deposition for a streamlined chitin production for multiple applications.

In insects, chitin is found in ectodermal tissues, where it forms chito-protein cuticles, and in the gut, where it forms a Peritrophic Matrix. Chitin is deposited to the extracellular space by Chitin Synthases (CHS) enzymes, which belong to the family of β-glycosyltransferases, that also includes Cellulose Synthases (CES) and Hyaluronane Synthases (HS). Most insect species encode two CHS types, CHS-A, required for chitin deposition in epidermis, trachea, foregut and hindgut, and CHS-B, required for chitin deposition in the midgut, as a principal component of the Peritrophic Matrix (Liu et al., 2019; Zhao, 2019; Zhu et al., 2016). The exact mechanism of chitin deposition is not fully elucidated yet, but it is proposed to occur in consecutive steps: 1) polymerisation by the catalytic domain of chitin synthase, 2) translocation through the chitin synthase of the nascent polymer across the membrane and release into the extracellular space, and 3) spontaneous assembly of translocated polymers to form crystalline microfibrils (Merzendorfer, 2006; Merzendorfer, 2011; Merzendorfer and Zimoch, 2003; Zhao, 2019; Zhu et al., 2016)

In *Drosophila*, CHS-A is encoded by *krotzkopf verkehrt* (*kkv*), which is responsible for chitin deposition in ectodermal tissues (Moussian et al., 2005; Ostrowski et al., 2002). But besides *kkv*, our previous work identified another function exerted by *expansion* (*exp*) and *rebuf* (*reb*) to be required for chitin deposition. Exp and Reb are two homologous proteins, containing a conserved Nα-MH2 domain, that serve the same function, as the presence of only one of them can promote chitin deposition. In the absence of *exp/reb*, no chitin is deposited in ectodermal tissues, in spite of the presence of *kkv*, indicating that this function is absolutely required. But most importantly, we found that *kkv* and *exp/reb* compose the minimal genetic network which is not only required, but also sufficient for chitin deposition. Thus, the concomitant expression of the two activities, *kkv*+*exp/reb*, promotes increased chitin deposition in ectodermally-derived tissues that normally deposit chitin, like the trachea, and ectopic chitin deposition in ectodermally-derived tissues that normally do not deposit chitin, like the salivary glands (Moussian et al., 2015). In spite of the capital importance of the *exp/reb* function in chitin deposition, the mechanism of activity of *exp/reb* has not been identified yet, nor their putative relation/interactions with *kkv*.

In this work we have investigated the cellular and molecular mechanisms of chitin deposition in *Drosophila* and the roles of *exp/reb* and *kkv* in the process. We have found that the activities of Kkv in chitin polymerisation and chitin translocation are uncoupled, and we propose that chitin translocation, but not chitin polymerisation, requires Exp/Reb activity. Our cellular analysis has revealed that when extracellular chitin deposition is prevented, Kkv-polymerised chitin accumulates in the cytoplasm as membrane-less punctae. In addition, we detected a clear correlation between Kkv function in chitin polymerisation and/or translocation and Kkv subcellular localisation. A molecular analysis of Exp/Reb and Kkv proteins, using a structure-function approach, revealed key functions of different conserved motifs of these proteins in chitin polymerisation and extracellular deposition and in protein subcellular localisation. A detailed analysis of the subcellular localisation of Exp/Reb and Kkv indicates that these proteins display a largely complementary pattern at the apical membrane. Furthermore, we find that Exp/Reb regulates the pattern of distribution of Kkv protein at the apical membrane. Based on the current understanding of the activity of glycosyltransferases like cellulose synthases and on the knowledge of *Drosophila* chitin synthase activity, we propose a model in which Exp/Reb regulate chitin deposition by modulating the distribution and organisation of Kkv complexes at the apical membrane, which would regulate the capacity of Kkv to translocate and release chitin fibers extracellularly.

## RESULTS

### 1. The activities of Kkv in chitin polymerisation and translocation are uncoupled and Exp/Reb activity is specifically required for chitin translocation

Chitin deposition is proposed to occur in 3 consecutive steps: 1) chitin polymerisation by CHS, 2) translocation of the nascent polymer through a CHS chitin-translocating channel across the membrane and release into the extracellular space, and 3) spontaneous assembly of translocated polymers to form crystalline microfibrils. In addition, it has also been proposed that the chitin polymerisation and translocation steps are tightly coupled (Merzendorfer, 2006; Merzendorfer, 2011; Merzendorfer and Zimoch, 2003; Zhao, 2019; Zhu et al., 2016). However, no experimental data is available on Kkv with respect to this model. We aimed to investigate this model by carrying out a molecular and cellular analysis of the roles of Kkv and Exp/Reb in chitin deposition.

We found that Kkv can promote extracellular chitin deposition (which requires chitin translocation) in ectodermal tissues only in combination with Exp/Reb activity. *kkv* overexpression in the tracheal system does not affect extracellular chitin deposition (which starts from stage 13 as in the wild type) or tracheal morphogenesis (Fig. 1A,B, S1A,C (Moussian et al., 2015). However, we detected the presence of intracellular chitin punctae at early stages (before stage 14) (Fig 1A’, S3), which indicates the ability of *kkv* to polymerise chitin. These intracellular chitin punctae disappeared from stage 14, when chitin is then deposited extracellularly (Fig 1B’). This switch from intracellular chitin to extracellular chitin perfectly correlates with the expression of *exp/reb* (Moussian et al., 2015), suggesting that *exp/reb* promote extracellular chitin deposition. In agreement with this, in *exp reb* mutants overexpressing *kkv*, we find intracellular chitin punctae until late stages and no extracellular chitin deposition (Fig 1C, S3, (Moussian et al., 2015)). We also find intracellular chitin punctae and no extracellular chitin when we expressed *kkv* in salivary glands, which do not express *exp* and *reb* (Fig. 1D, S3, (Moussian et al., 2015)). In contrast, the concomitant overexpression of *kkv* and *exp/reb* anticipates and increases extracellular chitin deposition in the trachea, which lead to tracheal morphogenetic defects (Fig S1A-D, S3 (Moussian et al., 2015)). In addition, *kkv* and *exp/reb* co-expression in salivary glands promotes chitin deposition in the lumen (Fig S1E, S3 and (Moussian et al., 2015). No intracellular punctae of chitin were detected under these conditions, suggesting that all chitin synthesised by Kkv is deposited extracellularly by Exp/Reb activity. Thus, we propose that the functions of Kkv in chitin polymerisation and translocation are uncoupled, and that Exp/Reb activity is required for chitin translocation and release to the extracellular space. In this context, the presence of intracellular chitin would reflect the activity of Kkv in chitin polymerisation that cannot be further processed and translocated due to the absence of Exp/Reb activity.

**Figure 1.**
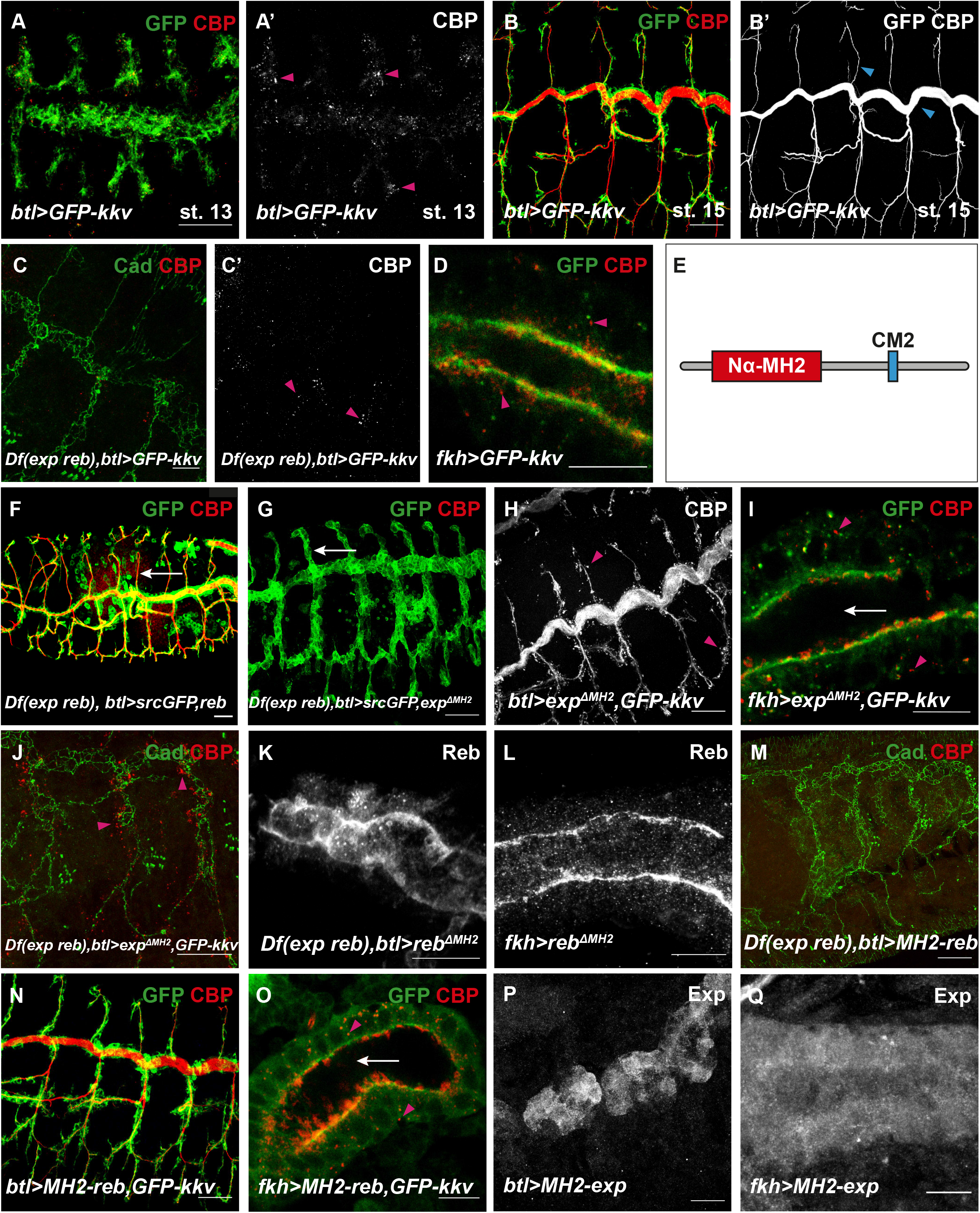
Analysis of the role of the Nα-MH2 domain of Exp/Reb. All images show projections of confocal sections, except D, I, L and O, which show single confocal sections. (A,B) The overexpression of *GFP-kkv* in the trachea leads to the presence of intracellular chitin vesicles at early stages (pink arrowheads) (A, A’). At later stages, intracellular chitin vesicles are not present and chitin is deposited extracellularly in the lumen (blue arrowheads) (B, B’). (C, C’) In *exp reb* mutants, the overexpression of *GFP-kkv* in the trachea produces intracellular chitin punctae until late stages (pink arrowheads). (D) The overexpression of *GFP-kkv* in salivary glands produces intracellular chitin vesicles (pink arrowheads). (E) Schematic representation of Exp protein. (F,G) In *exp reb* mutants, the expression of a wild type form of *exp/reb* rescues the lack of extracellular chitin deposition (F, white arrow points to presence of CBP), while *exp*^*ΔMH2*^*/reb*^*ΔMH2*^ do not (G, white arrow points to absence of CBP). (H,I) The co-overexpression of *GFP-kkv* and *exp*^*ΔMH2*^ in control embryo does not produce morphogenetic defects in trachea (H) or extracellular chitin diposition in salivary glands (white arrow) (I), however, intracellular chitin punctae are present (pink arrowheads in H,I). (J) The co-expression of *GFP-kkv* and *exp*^*ΔMH2*^ in *exp reb* mutants produces intracellular chitin particles (pink arrowheads) but does not rescue the lack of extracellular chitin deposition. (K, L) Reb^ΔMH2^ localises apically in trachea (K) and in salivary glands (L). (M) *MH2-reb* is not able to rescue the absence of extracellular chitin deposition in *exp reb* mutants. (N,O) The simultaneous expression of *MH2-reb* and *GFP-kkv* does not produce morphogenetic defects or ectopic chitin deposition in trachea (N) and in salivary glands (white arrow in O), but intracellular chitin vesicles are present (pink arrowhead in O). (P,Q) MH2-Exp protein does not localize apically in trachea (P) or in salivary gland (Q). Scale bars A-C,F-H,J,M,N 25 μm, D,I,K,L,O-Q 10 μm.

### 2. Structure-Function analysis of the roles of Exp/Reb in chitin translocation

We aimed to understand how Exp/Reb may regulate Kkv-dependent chitin translocation. The only recognisable domain of Exp/Reb identified was an Nα-MH2 (Beich-Frandsen et al., 2015; Moussian et al., 2015). However, in the course of this work, we identified a second domain highly conserved, which we called conserved motif 2 (CM2) (Fig 1E). We investigated the functional requirements of each of these two domains in chitin deposition. We generated different UAS transgenic mutant lines with the aim to evaluate their ability to rescue the lack of *exp reb* activity and their ability to promote chitin deposition when co-expressed with *kkv*. In agreement with their interchangeable activities, the results we obtained for *exp* or *reb* were comparable, so we will refer to them indistinctly.

#### 2.1. Nα-MH2 domain is required for chitin translocation

We generated Exp and Reb mutant proteins that lacked the Nα-MH2 domain (*exp*^*ΔMH2*^, *reb*^*ΔMH2*^). The expression of full length *exp* and *reb* rescues the lack of chitin deposition in the extracellular space in the trachea of *exp reb* mutants (Fig 1F, S3, (Moussian et al., 2015). In contrast, no extracellular chitin was deposited in a *exp reb* mutant background expressing *exp*^*ΔMH2*^ or *reb*^*ΔMH2*^ (Fig 1G), indicating that these proteins are not functional.

In agreement with a role for the Nα-MH2 in promoting extracellular chitin deposition, we found that co-expression of *kkv* and *exp*^*ΔMH2*^/*reb*^*ΔMH2*^ did not lead to tracheal morphogenetic defects (Fig 1H, S3), or ectopic chitin deposition in the lumen of salivary glands (Fig 1I), as full length *exp/reb* do (Fig S1D,E, S3 (Moussian et al., 2015)). However, as expected, the lack of extracellular chitin deposition was accompanied by the presence of intracellular chitin punctae (Fig 1I,J, S3). These results indicated that the activity of Exp/Reb in extracellular chitin deposition resides in its Nα-MH2, suggesting a role for the Nα-MH2 domain of Exp/Reb in chitin translocation and release to the extracellular space.

To discard an unspecific effect on extracellular chitin deposition due to the absence of the MH2 domain, we investigated Exp^ΔMH2^/Reb^ΔMH2^ localisation. We found that Reb^ΔMH2^ localised apically when expressed in the trachea (Fig 1K) or salivary glands (Fig 1L), in a comparable pattern to the overexpression of full length Reb protein (Fig S2A,B). These results indicated that the Nα-MH2 domain is not required for Exp/Reb localisation and instead plays a specific role in extracellular chitin deposition.

As the Nα-MH2 is required for chitin extracellular deposition, we asked whether it was also sufficient to promote extracellular chitin deposition. We generated UAS constructs with only the Nα-MH2 domain of Exp and Reb (*MH2-exp* and *MH2-reb*). We found that the constructs were unable to rescue the lack of extracellular chitin deposition in *exp reb* mutants and to produce tracheal morphogenetic defects in the trachea or chitin luminal disposition in salivary glands when co-expressed with *kkv* (Fig 1M-O, S3). This indicated that the MH2-Exp/MH2-Reb proteins are not functional to promote extracellular chitin deposition. In agreement with this observation, we also detected the presence of numerous intracellular puncta of chitin (Fig 1O, S2C, S3). Correlating with this lack of activity, we observed that MH2-Exp did not localise apically, as Exp does (FigS2D), and instead it was found in the cytoplasm in the trachea or salivary glands (Fig 1P,Q). Our results suggested that either other domains of the protein also contribute to extracellular chitin deposition, or that apical localisation is required for Exp/Reb activity. The results also indicated that other domains in Exp/Reb are required for apical localisation.

In summary, the analysis of the Nα-MH2 domain points that it is required for translocation and extracellular release of chitin but it is dispensable for protein localisation and chitin polymerisation.

#### 2.2. The Conserved Motif 2 is required for Exp localisation

We searched for conserved domains by comparing the amino acid sequences of several Exp homologs. We identified a highly conserved region not previously described in the literature. The Conserved Motif 2 (from now on CM2) contained a region of 8 aa highly conserved (blue square in Fig 2A) followed by a region of 9 aa less conserved (red square in Fig 2A). To investigate the functional activity of the CM2 we generated *UASexp*^*ΔCM2*^ transgenic lines.

**Figure 2.**
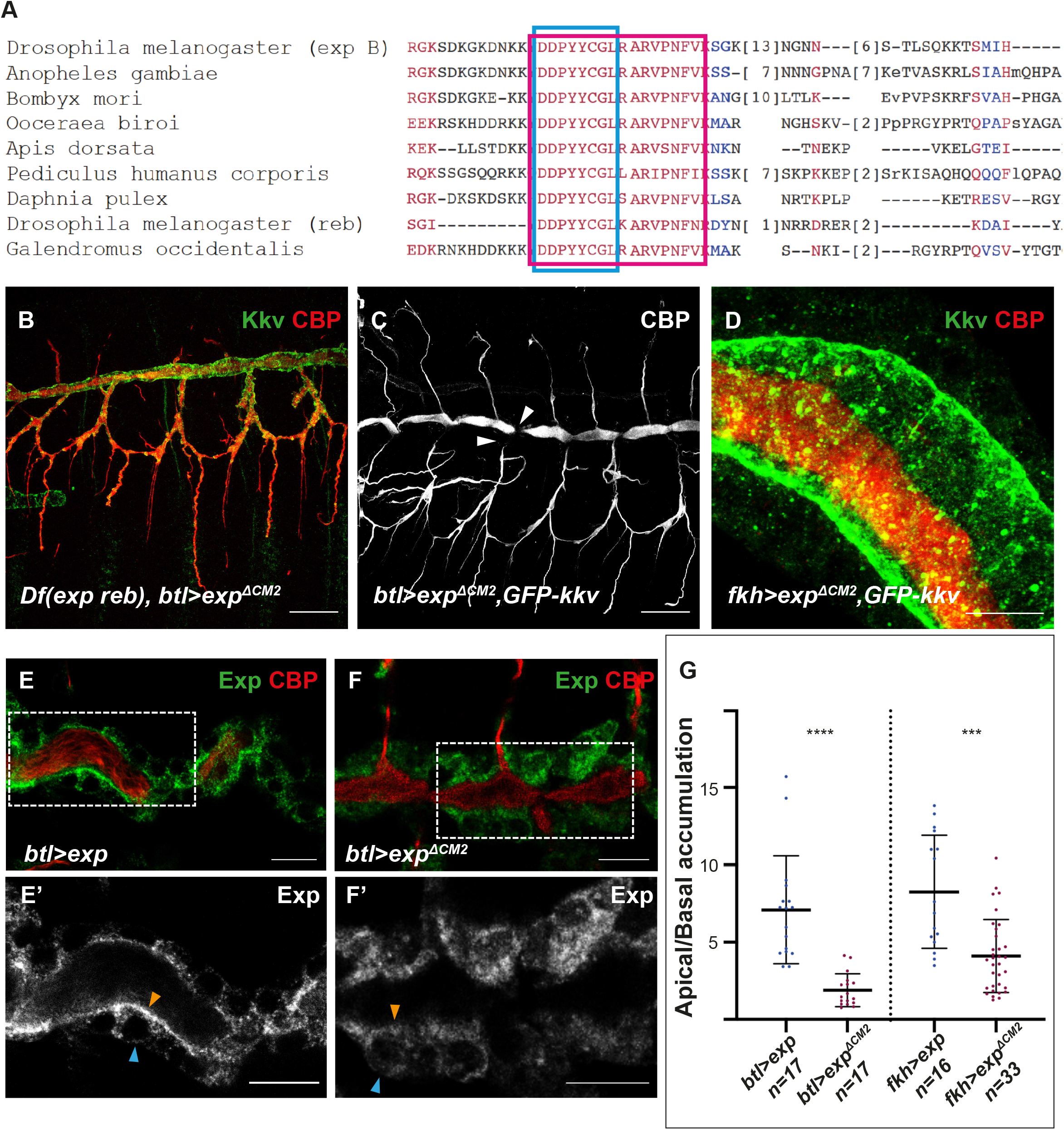
Analysis of the of the role of the CM2 domain of Exp/Reb. (A) Alignment of amino acids (aa) sequences of the isoform B of Exp (aa 356-433) and homologs; the blue square indicates 8 aa highly conserved, the red square includes 9 aa less conserved. (B-D) Show projections of confocal sections and (E-F) show single confocal sections. (B) The overexpression of *exp*^*ΔCM2*^ in a *exp reb* mutant background rescues the lack of extracellular chitin deposition. (C,D) The simultaneous expression of *exp*^*ΔCM2*^ and *GFP-kkv* produces morphogenetic defects in the trachea (arrowheads in C) and ectopic chitin deposition in the lumen of salivary glands (D). (E,F) Overexpressed Exp localises mainly in the apical region (orange arrowheads) with respect the basal domain (blue arrowheads), while the apical accumulation of overexpressed Exp^ΔCM2^ is less conspicuous (F). (G) Quantifications of accumulation of Exp and Exp^ΔCM2^ in apical versus basal region. Scale bars B,C 25 μm, D-F 10 μm.

The expression of *exp*^*ΔCM2*^ in a *exp reb* mutant background rescued the lack of extracellular chitin deposition (Fig 2B, S3), indicating that the protein is functional. In agreement with this, when co-expressed with *kkv* in the trachea, it produced morphogenetic defects (Fig 2C, S3), comparable to the overexpression of *kkv* and *exp/reb* (Fig S1D, S3). Similarly, expression of *kkv* and *exp*^*ΔCM2*^ in salivary glands produced ectopic chitin deposition in the luminal space (Fig 2D). These results indicated that the CM2 is not required for chitin polymerisation and translocation to the extracellular space. In agreement with no requirements of the CM2 in chitin translocation, no intracellular chitin punctae were detected when co-expressing *kkv* and *exp*^*ΔCM2*^ (Fig 2D, S3).

To further investigate the role of the CM2, we analysed protein localisation. The endogenous Exp and Reb proteins localise mainly apically at the membrane, although a bit of the protein can be detected intracellularly (Fig S2D and (Moussian et al., 2015)). This pattern of subcellular accumulation was maintained when overexpressing Exp (Fig 2E). In contrast, overexpressed Exp^ΔCM2^ did not show such a distinct apical localisation (Fig 2F). We analysed the ratio of accumulation of the Exp proteins (control and Exp^ΔCM2^) in apical versus basal regions. Quantifications indicated a decreased apical enrichment of Exp^ΔCM2^ compared to full length Exp (Fig. 2H). This result indicated that the CM2 is involved in Exp/Reb localisation. However, we could still detect some accumulation of Exp^ΔCM2^ at the apical membrane (Fig 2F), which correlated with its activity in chitin deposition.

In summary, we identified a highly conserved domain in Exp proteins that is dispensable for chitin polymerisation and translocation but is required for correct protein localisation, which we propose is important for Exp/Reb activity (see discussion).

### 3. Structure-Function analysis of two conserved Kkv domains in chitin deposition

Kkv is a large protein with multiple functional domains, and several transmembrane domains (Adler, 2019; Merzendorfer, 2011; Moussian, 2013). As for other members of the β-glycosyltransferase family, it is proposed that the activity of CHS, like Kkv, depends on oligomerisation and interactions with other proteins (Merzendorfer, 2011; Zhao, 2019; Zhu et al., 2016). We investigated two Kkv domains with putative roles in direct or indirect interactions with other proteins or in oligomerisation (Moussian, 2013; Moussian et al., 2005), the conserved motif WGTRE (aa 1076-1080) and a coiled-coil domain (aa 1087-1107) (Fig 3A).

**Figure 3.**
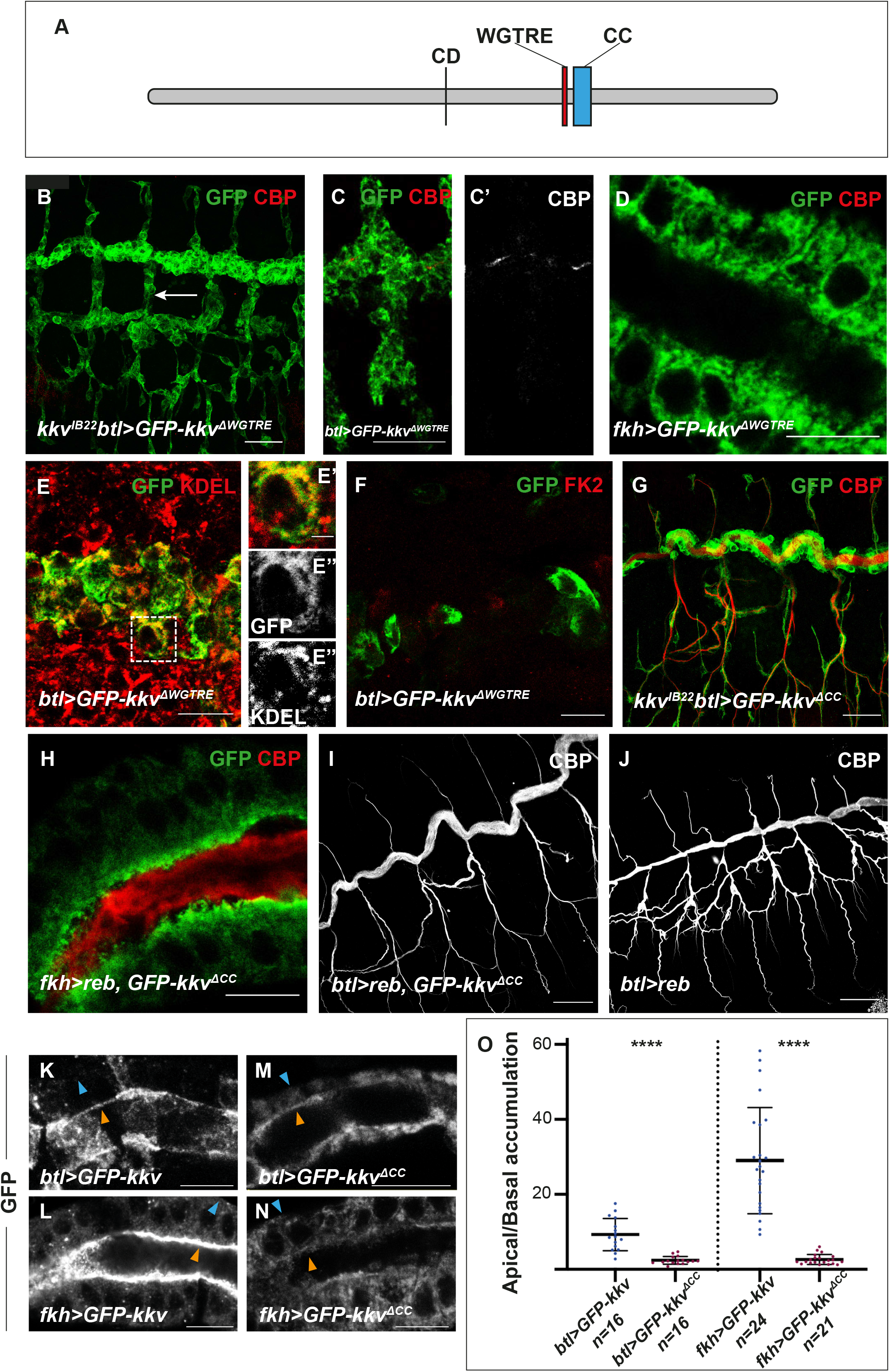
Analysis of the WGTRE and CC domains of Kkv. (A) Schematic representation of Kkv protein (CD, Catalytic Domain; WGTRE; CC, Coiled-Coil domain). (B,C,G,I,J) Show projections of confocal sections and (D-F,H,K-N) show single confocal sections. (B) The overexpression of *GFP-kkv*^*ΔWGTRE*^ in a *kkv* mutant background does not rescue the absence of extracellular chitin deposition (white arrow, note the absence of CBP) and the protein accumulates in a generalised pattern. (C-D) The overexpression of *GFP-Kkv*^*ΔWGTRE*^ does not produce intracellular chitin punctae, neither in trachea at early stages (C-C’), nor in salivary glands (D). (E-E’’’) GFP-kkv^ΔWGTRE^ colocalise with the ER marker KDEL. (F) GFP-Kkv^ΔWGTRE^ does not colocalise with the marker FK2. (G) The overexpression of *GFP-kkv*^*ΔCC*^ in a *kkv* mutant background rescues the lack of extracellular chitin deposition in the trachea (note the presence of CBP staining). (H,I) The simultaneous expression of *reb* and *GFP-kkv*^*ΔCC*^ in salivary glands produces ectopic extracellular chitin (H), and no defects in trachea (I). (J) The overexpression of *reb* in trachea leads to morphogenetic defects. (K,L) Overexpressed GFP-Kkv localises mainly apically (orange arrowheads) although a bit of the protein can be detected in the basal region (blue arrowheads). (M,N) Apical accumulation of overexpressed GFP-Kkv^ΔCC^ is less conspicuous. (O) Quantifications of accumulation of GFP-Kkv and GFP-Kkv^ΔCC^ in apical versus basal region. Scale bars B,C,G,I,J 25 μm, D-F,H,K-N 10 μm.

#### 3.1. The WGTRE domain is required for Kkv ER-exit

The conserved motif WGTRE was proposed to be an essential domain for Kkv activity as a point mutation changing the glycine renders an inactive protein (Ostrowski et al., 2002). This domain has been suggested to be involved in oligomerisation or interactions with other factors (Moussian et al., 2005)). We generated a protein lacking this domain, GFP-Kkv^ΔWGTRE^.

To determine the activity of this protein we assayed its rescuing capacity in a *kkv* mutant background. While a wild type form of *kkv* can rescue the absence of chitin produced by the absence of *kkv* (Moussian et al., 2015), *GFP-kkv*^*ΔWGTRE*^ could not, and trachea was defective (Fig 3B, S3). This indicated that the protein is not functional, confirming that the WGTRE domain is essential for chitin production. Accordingly, expression of *GFP-kkv*^*ΔWGTRE*^ did not produce chitin vesicles in the trachea at early stages (Fig 3C, S3), or in salivary glands (Fig 3D), as GFP-kkv does (Fig 1A,D, S3), indicating absence of chitin polymerisation..

To better understand the role of this domain we analysed the localisation of the GFP-Kkv^ΔWGTRE^ protein. We found no apical accumulation of this protein, neither in a wild type background nor in a *kkv* mutant background (Fig 3B). Instead, we found a generalised pattern in the cytoplasm. Co-stainings with the ER marker KDEL (Fig 3E) indicated that the GFP-Kkv^ΔWGTRE^ protein is retained in the ER.

ER retention may be due either to a defective folding of the protein or to a specific effect of this domain in Kkv trafficking to the membrane. Proteins with a defective folding are degraded from ER upon ubiquitination (Tsai and Weissman, 2010). To distinguish between these two possibilities, we used the FK2 antibody that recognises mono-and polyubiquitinated conjugates, but not free ubiquitin (Fujimuro et al., 1994). From our results, GFP-Kkv^ΔWGTRE^ and FK2 do not colocalise (Fig 3F), indicating that the protein is not ubiquinated. The results strongly suggested a role for the WGTRE domain in Kkv trafficking from the ER to the membrane. ER exit is required for chitin polymerisation by Kkv

#### 3.2. The coiled-coil domain is required for Kkv localisation and full Kkv activity

Class A chitin synthases contain a coiled-coil domain localised C-terminal to the active centre. Potentially, the coiled-coil domain could mediate association to yet unknown partner/s, or be involved in protein oligomerisation, regulating chitin synthase localisation or activity (Adler, 2019; Merzendorfer, 2006; Moussian, 2013).

We generated a protein lacking the coiled-coil region, GFP-Kkv^ΔCC^. This mutant protein rescued the lack of chitin in a *kkv* mutant background (Fig 3G, S3), indicating that it is functional. Accordingly, the concomitant expression of *reb* and *GFP-kkv*^*ΔCC*^ in salivary glands resulted in deposition of chitin in the luminal space (Fig 3H, S3). Altogether these results indicated that GFP-Kkv^ΔCC^ acts as a functional protein in these contexts.

We found, however, a condition in which *GFP-kkv*^*ΔCC*^ behaved differently from *GFP-kkv*. The overexpression of *reb* and full length *GFP-kkv* in the trachea produced strong morphogenetic defects (Fig S1D, S3 and (Moussian et al., 2015)). In contrast, the overexpression of *reb* and *GFP-kkv*^*ΔCC*^ did not produce this abnormal phenotype, and instead tubes and chitin deposition appeared normal (Fig 3I, S3). Importantly, the overexpression of *reb* alone also leads to morphogenetic defects, due to the presence of endogenous *kkv* (Fig 3J, S3). Because the overexpression of *reb* and *GFP-kkv*^*ΔCC*^ (in the presence of endogenous *kkv*) reverts the tracheal defects of overexpression of *reb* alone, our results suggest that the *GFP-kkv*^*ΔCC*^ is interfering with the endogenous wild type kkv, which can no longer produce a dominant effect in combination with extra Reb.

We analysed the localisation of GFP-Kkv^ΔCC^ to further determine the roles of the CC domain. We found that the protein can still localise at the apical domain, as GFP-Kkv does (Fig 3K-N), however, we found that the apical enrichment was not as clear as for GFP-Kkv. Quantification of the ratio of protein accumulation in the apical domain versus the basal domain indicated significant differences with respect the control. We confirmed these observations in salivary glands (Fig 3O).

Altogether these results indicate that the coiled-coil domain is dispensable for polymerisation and translocation of chitin, but plays a role in protein localisation. In addition, the results suggest that GFP-Kkv^ΔCC^ can interfere with endogenous Kkv, which may indicate a role in protein oligomerisation (see discussion).

### 4. Cellular analysis of chitin polymerisation and extracellular translocation

We aimed to investigate further at the cellular level the roles of Kkv in chitin polymerisation and translocation and the nature of the intracellular chitin deposition. To this purpose we used the salivary glands. As shown previously, the co-expression of *kkv* and *exp/reb* in salivary glands (which in normal conditions do not produce chitin) leads to the deposition of chitin extracellularly in the lumen (Fig 4A, S1E and (Moussian et al., 2015). In contrast, in the absence of *exp/reb* activity, like when co-expressing *kkv* and *exp*^*ΔMH2*^*/reb*^*ΔMH2*^, we find intracellular chitin accumulation and no extracellular chitin in the lumen (Fig 4B,C). As indicated before, this suggested that chitin is deposited intracellularly because it cannot be translocated to the extracellular space. However, other mechanisms could explain intracellular chitin accumulation. For instance, intracellular accumulation could result from an abnormal endocytic uptake of previously translocated chitin occurring in the absence of *exp/reb* activity. To test this possibility, we blocked endocytosis in *GFP-kkv+exp*^*ΔMH2*^*/reb*^*ΔMH2*^ expressing conditions. We still detected intracellular chitin punctae, indicating that these result from lack of chitin translocation (Fig 4D).

**Figure 4.**
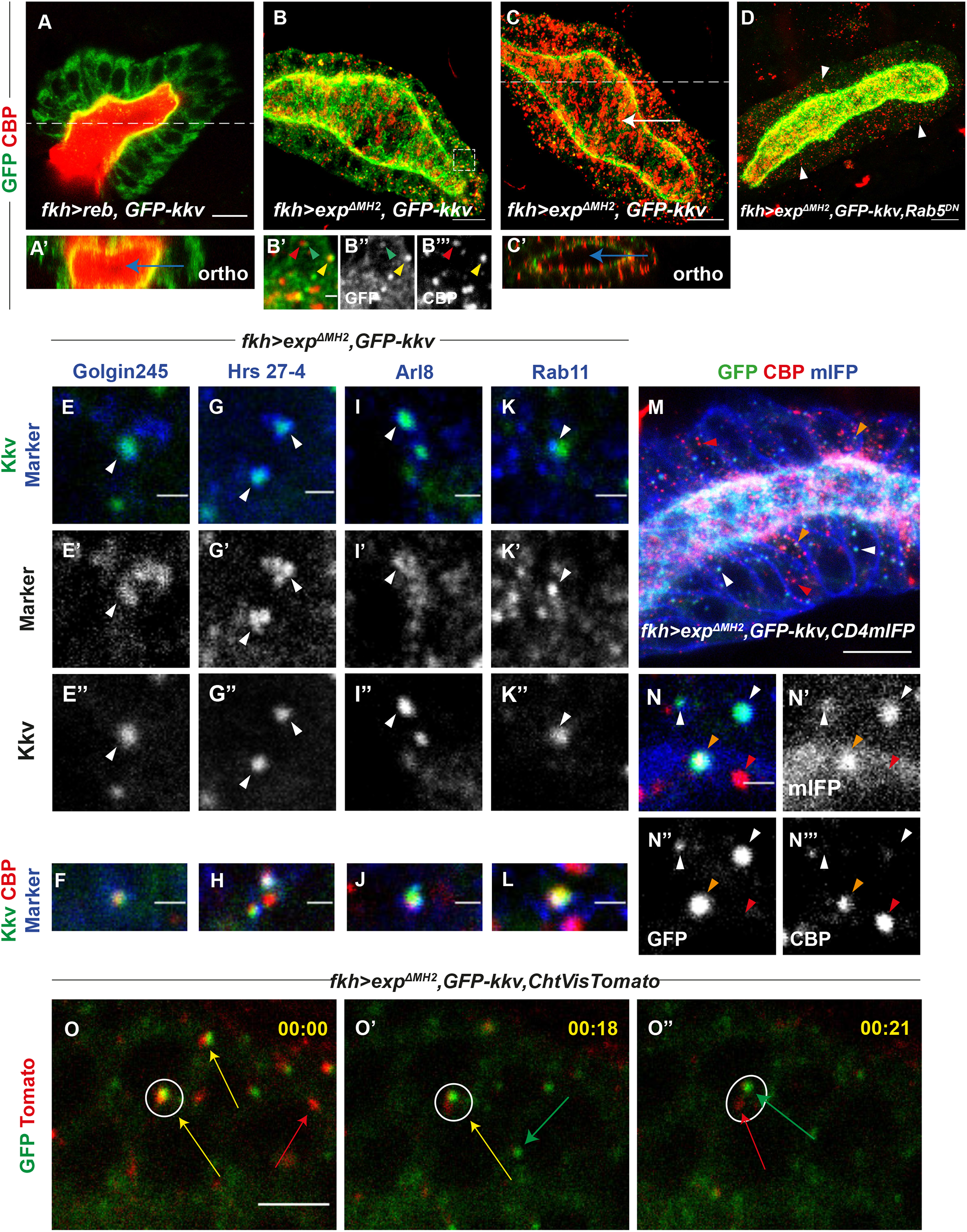
Analysis of intracellular chitin deposition. All images show salivary glands. (A,E-L,N,O) Show single confocal sections and (B-D,M) show projections of several sections. (A-A’) The concomitant expression of *reb* and *GFP-kkv* leads to luminal chitin deposition (blue arrow in orthogonal section in A’). (B,C) Co-expression of *exp*^*ΔMH2*^ and *GFP-kkv* produces intracellular chitin punctae, some of which colocalise with GFP-Kkv vesicles (yellow arrowhead) while others do not (red arrowhead). GFP-Kkv vesicles without chitin are also observed (green arrowhead). Note the accumulation of chitin in the apical domain (white arrow in C) that is not deposited extracellularly in the lumen (blue arrow in orthogonal section in C’) (D) In *Rab5*^*DN*^ background, intracellular chitin punctae are still present (white arrowheads). (E-L) Analysis of the nature of GFP-Kkv vesicles and chitin punctae using markers Golgin245 (E-F), Hrs 27-4 (G-H), Arl8 (I-J) and Rab11 (K-L); arrowheads indicate colocalisation between Kkv and each specific marker. (M,N) All GFP-Kkv vesicles colocalise with the membrane marker CD4-mIFP (white arrowheads) and few of them contain also chitin (orange arrowheads); single chitin punctae do not colocalise with CD4-mIFP (red arrowheads). (O-O’’) Frames from live imaging movie show that common GFP-Kkv and chitin punctae (yellow arrow) can separate from each other; however, many GFP-Kkv (green arrow) and chitin puncta (red arrow) do not colocalise. Scale bars A-D,M 10 μm, E-L, N-N’’’ 1 μm, O-O’’ 5 μm.

The use of the *UAS-GFP-kkv* line served us to overexpress *kkv*, but also to visualise *kkv* accumulation. Expression of *GFP-kkv* in salivary glands revealed the expected Kkv localisation at the apical membrane, but also the presence of many intracellular Kkv punctae in the cytoplasm. We asked whether intracellular chitin punctae and Kkv punctae colocalised. We found that around a 34% of GFP-Kkv punctae contained chitin, and around 20% of intracellular chitin punctae were also positive for GFP-Kkv (Fig 4B). We also found many examples in which GFP-Kkv and chitin punctae were in very close proximity (Fig 4B-B’’). Intracellular chitin punctae distributed throughout the cytoplasm, but we also detected large amounts of chitin deposits at the apical domain of the cell (Fig 4C). Nevertyheless, orthogonal sections showed that this apical chitin is not deposited extracellularly in the lumen (Fig 4C’), but instead localised intracellularly in the apical region, in contrast to the co-expression GFP-Kkv and wild type Reb (Fig 4A’).

To identify the nature of Kkv and chitin punctae, we performed colocalisation analysis with different markers of intracellular trafficking. Colocalisation with the TransGolgi marker Golgin 245 indicated that several GFP-Kkv punctae corresponded to secretion vesicles (around 12% of GFP-Kkv vesicles) (Fig 4E). A few of these GFP-Kkv secretion vesicles contained chitin (Fig 4F). We also detected that many GFP-Kkv vesicles colocalised with late endosomal markers (around 60% of GFP-Kkv vesicles), like the ESCRT-0 complex component Hrs, indicating an endocytic recycling of Kkv (Fig 4G). Some of these endocytic Kkv vesicles also contained chitin (Fig 4H). To investigate whether endocytosed Kkv was then following the degradation pathway or was recycled back to the membrane we used lysosomal markers and markers for vesicle recycling. We found colocalisation of GFP-Kkv and the lysosomal marker Arl8 (35% of GFP-Kkv vesicles), indicating that part of the protein is degraded (Fig 4I,J). Finally, we also found colocalisation of GFP-Kkv with Rab11 (around 15% of GFP-Kkv vesicles), a marker for recycling (Fig 4K). Again, a few of them also contained chitin (Fig 4L). Altogether the results suggested that Kkv is transported to the membrane, internalised, and recycled back to the membrane or degraded. During this trafficking route, Kkv protein is able to polymerise chitin.

We realised that many intracellular chitin punctae did not colocalise with GFP-Kkv or with any other trafficking marker, suggesting that they may not correspond to intracellular vesicles. To determine whether these chitin punctae accumulated in membrane-confined compartments we used the general plasma membrane marker CD4::mIFP (Yu et al., 2015), which is enriched in plasma membrane and other subcellular membrane compartments (Fig 4M). We found that while all GFP-Kkv vesicles (both those containing and non-containing chitin) colocalised with the CD4::mIFP marker, the single chitin punctae did not (Fig 4N). This result suggests that these intracellular chitin punctae correspond to short chitin fibers floating freely in the cytoplasm.

To further understand the nature and dynamics of these chitin punctae, we performed live imaging using a live-probe for chitin and GFP-Kkv. The results confirmed that many GFP-Kkv and chitin punctae did not colocalise. Nevertheless, we found examples in which we detected common punctae of GFP-Kkv and chitin, and observed that KkvGFP and chitin were then separating from each other (Fig 4O, S4, Movies 1,2).

Altogether our results are consistent with a model in which Kkv can polymerise chitin in a constitutive manner, and when this chitin is not extruded to the extracellular domain it accumulates intracellularly as short fibers.

### 5. Kkv activity correlates with Kkv trafficking and localisation

Previous experiments suggested that Kkv activity plays a role in Kkv localisation (Adler, 2019). In wild type embryos, endogenous Kkv protein (visualised with anti-Kkv) (Fig 5A) is found localised apically and in intracellular vesicles (Fig 5A), as it is the case of GFP-Kkv expressed in salivary glands. We investigated whether this pattern of localisation was affected by Kkv activity.

**Figure 5.**
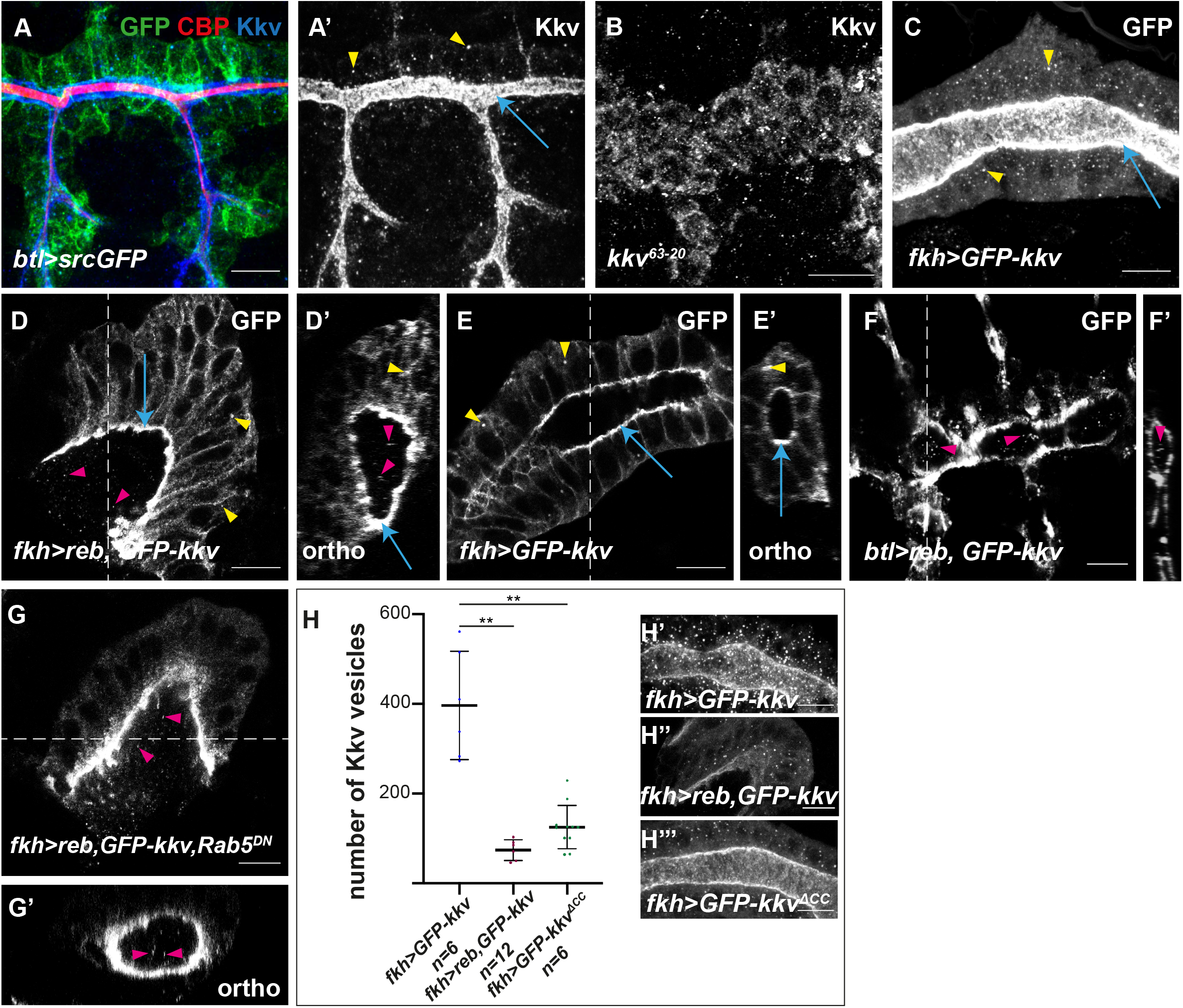
Analysis of Kkv trafficking. All images are single confocal sections except A-C, which are projections of confocal sections. (A,A’) In trachea of wild type embryos, Kkv is present in the apical region (blue arrows) and in intracellular vesicles (yellow arrowheads). (B) In *kkv* mutants unable to polymerise chitin, Kkv is not properly localised. (C) GFP-Kkv localises to the apical region (blue arrows) and in intracellular vesicles (yellow arrowheads). (D) When *reb* and *GFP-kkv* are co-expressed in salivary glands, Kkv is present in the apical membrane (blue arrow), in intracellular vesicles (yellow arrowhead) and also in punctae in the lumen (pink arrowheads). This is clearly observed in orthogonal sections (D’). (E,E’) In contrast, when *GFP-kkv* is expressed alone, luminal punctae are absent, and Kkv is only found apically (blue arrow) and in intracellular vesicles (yellow arrowhead). (F) Luminal punctae (pink arrowheads) are also observed in the trachea of embryos overexpressing *reb* and *GFP-kkv*. (G) When endocytosis is prevented, the co-expression of *reb* and *GFP-kkv* in salivary glands still leads to formation of Kkv luminal punctae (pink arrowheads). (H) Quantifications of the number of intracellular Kkv vesicles in salivary glands when expressing *GFP-kkv* (H’), *reb* and *GFP-kkv* (H’’) and *GFP-kkv*^*ΔCC*^. Scale bars 10 μm.

We found that in conditions of lack of chitin polymerisation (i.e. in amorphic *kkv* mutants with a point mutation in the catalytic domain, *kkv*^*63-20*^ mutants (Moussian et al., 2005)), Kkv did not localise apically and instead it was detected in the cytoplasm (Fig 5B). In conditions where chitin is polymerised but not translocated and accumulates in intracellular punctae (i.e. when expressing *GFP-kkv* in the absence of *exp/reb* activity), Kkv is secreted, reaches the membrane, is then internalised and degraded or recycled back to the membrane, as previously described (Fig 5C and Fig 4). In conditions where chitin is massively produced and deposited extracellularly (i.e. when expressing *reb* and *GFP-kkv* in salivary glands), we detected Kkv at the apical membrane and in vesicles (Fig 5D), although we detected less vesicles than with *GFP-kkv* alone (Fig 5C,H). In addition, we detected punctae of GFP-Kkv in the lumen of salivary glands (pink arrow in 5D), which were not detected in conditions of GFP-Kkv expression alone (Fig 5E). This extracellular punctae correlates with a huge production and extracellular deposition of chitin by *kkv*. In agreement with this observation, we also found GFP-Kkv punctae in the lumen of the trachea overexpressing *GFP-kkv* and *reb* (Fig 5F), which deposit increased amounts of chitin. These punctae seem similar to the extracellular Kkv punctae described in (Adler, 2019; Adler, 2020), and could therefore reflect Kkv shedding. Proteins can be shed to the extracellular space as extracellular vesicles, which comprise exosomes (derived from the endocytic trafficking) and microvesicles (directly shed from the plasma membrane) (Tricarico et al., 2017; van Niel et al., 2018). We could detect extracellular vesicles in salivary glands expressing *reb+GFP-kkv* in which we blocked the endocytic uptake, by expressing *Rab5*^*DN*^ (Fig 5G). This result suggested that the Kkv vesicles that we detect extracellularly may correspond to microvesicles (see discussion).

Finally, we also found differences in Kkv localisation and trafficking when the protein lacks the CC domain (Fig 5H). We observed a reduced number of GFP-Kkv^ΔCC^ vesicles compared to GFP-Kkv. This result suggests that the CC domain is directly or indirectly involved in Kkv trafficking.

Altogether our results point to a correlation between Kkv activity and Kkv trafficking and localisation.

### 6. Exp/Reb activity is required for Kkv apical distribution

Because we observed a correlation between Kkv activity and Kkv localisation, we asked whether *exp/reb* activity regulates Kkv localisation. We had previously shown that *exp/reb* activity is not required for apical localisation of overexpressed *GFP-kkv* (Moussian et al., 2015). However, the overexpression of *kkv* in this experimental setting could mask a possible role of *exp/reb* in Kkv localisation. Thus, we revisited this issue analysing the accumulation of endogenous Kkv using an antibody that we generated. We confirmed that Kkv localised apically in the absence of *exp/reb* (Fig 6A,B). We found that Kkv also localised apically when expressing the non-functional proteins Exp^ΔMH2^ or MH2-exp in trachea (Fig 6C,D).

**Figure 6.**
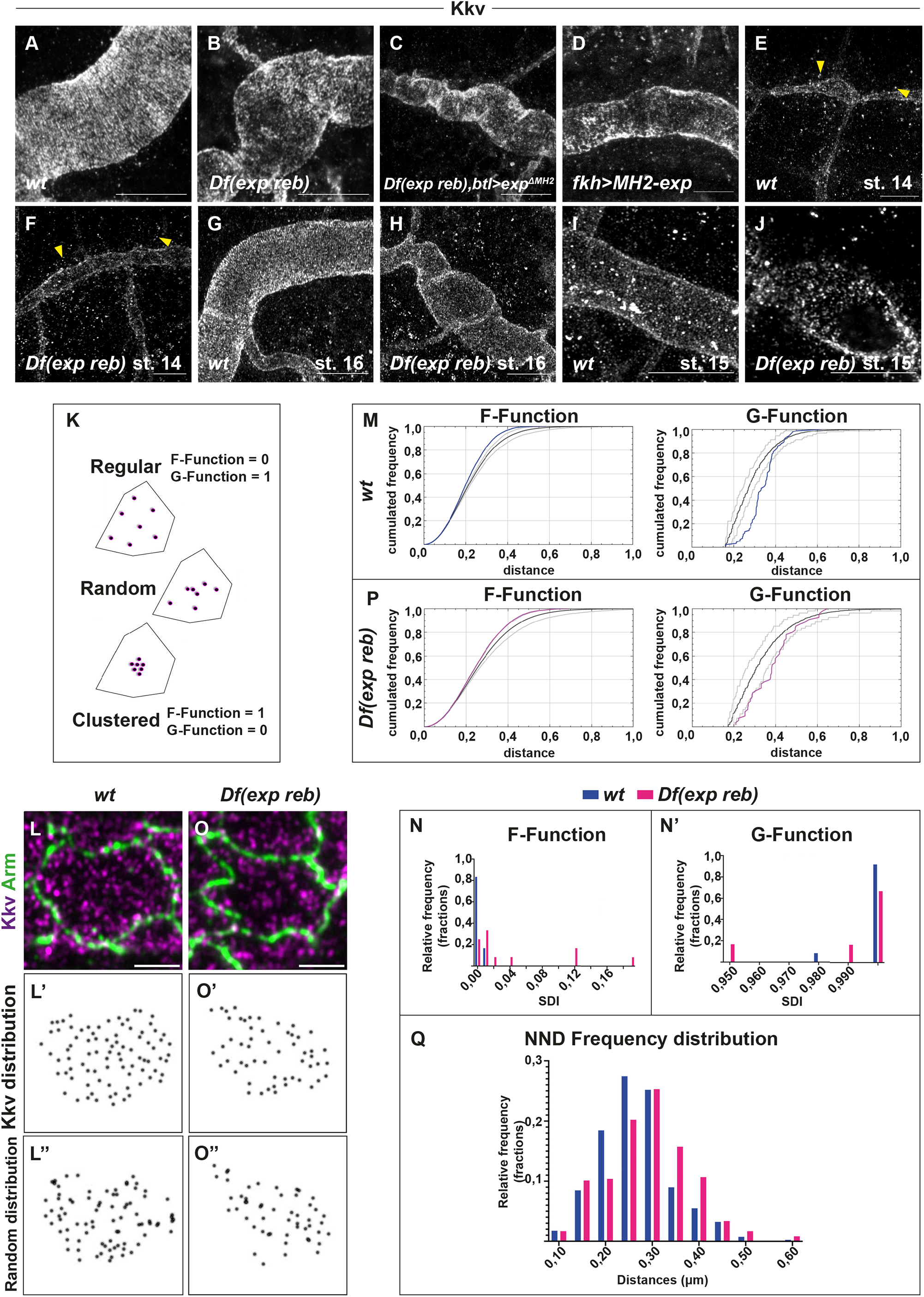
Analysis of Kkv apical distribution. All images are projections of confocal sections, of super-resolution microscopy. (A, B) Kkv localises apically in the trachea of wild type embryos (A) and in absence of *exp reb* (B). (C, D) The localisation of Kkv is apical also in presence of *exp* ^*ΔMH2*^ in trachea (C) and in presence of *MH2-exp* in salivary glands (D). (E, F) At stage 14, in wild type embryos (E) and in embryos deficient for *exp* and *reb* (F), Kkv is present in the apical membrane and in many intracellular vesicles (yellow arrowheads). (G) At stage 16, in wild type embryo, Kkv apical distribution follows the pattern of taenidial folds and intracellular vesicles are mostly absent. (H) At stage 16, in *exp reb* mutant embryos, Kkv is apical but it is not distributed uniformly. (I, J) At stage 15, in control embryos, Kkv pattern is apical, uniform and homogenous (I); instead in *exp reb* mutant embryos, Kkv distribution is not homogeneous at the apical membrane (J). (K) Three different types of spatial distribution within a selected area. The positions of the defined objects can be random, exhibit characteristics of attraction (clustered pattern) or repulsion (regular pattern). The F-Function, tends to be larger (≈1) for clustered patterns and smaller (≈ 0) for regular. The G-Function tends to be smaller (≈ 0) for clustered and larger (≈ 1) for regular patterns. (L) Kkv on the apical cell surface follows a regular distribution pattern. Kkv punctae (Magenta) on the apical cell area marked by Armadillo (Green) of a control embryo. (L’) Positions of Kkv punctae on the selected area marked by black dots. (L’’) Random pattern of distribution for the same area created by the spatial statistics 2D/3D image analysis plugin. (M) The corresponding observed F and G function (blue) are displayed above and below the reference simulated random distributions (black) and the 95% confidence interval (light gray) respectively, indicating a regular spatial pattern. (N) SDI histogram for the F-Function of the control (blue) and the *Df(exp reb)* samples. A significant difference between the frequency distributions for each group of individuals has been observed. (Kolmogorov-Smirnov D=0.5833, *p<0.05*) (N’) SDI histogram for the G-Function of the control (blue) and the *Df(exp reb)* samples. Statistical analysis of the distributions did not reveal significant differences between the two groups of individuals for this parameter (Kolmogorov-Smirnov D=0.25, *p>0.05*). (O) Kkv punctae (Magenta) on the apical cell area marked by Armadillo (Green) of a *exp reb* mutant embryo. (O’) Positions of Kkv punctae on the selected area marked by black dots. (O’’) Random pattern of distribution for the same area created by the spatial statistics 2D/3D image analysis plugin. (P) The corresponding observed F and G function (blue) are displayed above and below the reference simulated random distributions (black) respectively. Both curves largely overlap with the 95% confidence interval (light gray), indicating a tendency towards a random spatial pattern. (Q) Frequency distribution histograms for the Nearest Neighbour Distances between Kkv punctae in control (blue) and *exp reb* mutant samples. The distribution of values between the two groups is found significantly different (Kolmogorov-Smirnov D=0.2036, *p<0.005*). Scale bars A-J 10 μm. L,O 2μm

To investigate in detail possible subtle differences in Kkv apical accumulation, we used super-resolution microscopy and compared control embryos and *exp reb* mutants at different stages. At stage 14, both in control and *exp reb* mutant embryos, Kkv was found apical but also in many intracellular vesicles. Kkv at the apical membrane showed a non-uniform pattern in intensity and distribution (Fig 6E,F). At stage 16, control embryos showed a highly homogeneous apical distribution of Kkv in stripes, corresponding to the taenidial folds, and Kkv vesicles were largely absent (Fig 6G). In *exp reb* mutants at stage 16, Kkv was also apical, but in contrast to the control, Kkv did not distribute in lines and instead we observed a non-uniform pattern of distribution (Fig 6H). Because in *exp reb* mutant conditions chitin is not deposited and taenidia do not form (Moussian et al., 2015), this raised the possibility that the effects observed are due to the absence of taenidia. To discard this possibility, we analysed stage 15 embryos in which the chitinous cuticle forming the taenidia is not yet deposited but chitin is strongly deposited in the luminal filament. In control embryos we detected a very uniform and homogenous pattern of apical Kkv (Fig 6I). In *exp reb* mutants we observed a non-uniform distribution of apical Kkv protein with different intensities, resembling earlier stages (Fig 6J). To characterise further the topological distribution of Kkv on the apical area and identify possible alterations in *exp reb* mutants, we used spatial statistics to analyse our samples based on two main parameters; the distance between each Kkv punctum and its closest neighbour within a reference area defined by the apical marker Armadillo, and the distance between the puncta and arbitrary positions within the same reference area. Relevant to these parameters, two different cumulative distribution functions (CDF) defined as G and F - Function were calculated (Fig 6K), followed by the spatial distribution index (SDI), that accounts for the difference between the observed pattern and a completely random one (Andrey et al., 2010; Arpon et al., 2021; Ollion et al., 2013). For randomly organised patterns, the SDI within a population should be uniformly distributed between 0 and 1. Deviations from spatial randomness, are expected to shift the SDI values closer to 0 or 1, indicating regular (F–SDI=0, G–SDI=1) or clustered (F– SDI=1, G–SDI=0) organisation. The results of this analysis, showed that Kkv on the apical membrane, is evenly distributed following a regular pattern (Fig. 6L,L’,L’’,M). The SDI for the F-Function values for the control samples analysed were distributed between 0 - 0.01, while the one for the G-Function values were distributed between 0.98-1 (n=12) (Fig 6N,N’). The same analysis for the *exp reb* mutants (Fig 6O,O’,O’’,P), revealed low F and high G-function SDI values, within a range of 0 - 0.19 and 0.95 - 1 (n=12) respectively (Fig 6N,N’). Although this could also resemble a regular pattern within a population, in our comparisons the distribution of the F-Function SDI values of the controls was significantly different from the *exp reb* mutants (Kolmogorov-Smirnov D=0.5833, *p<0.05*) (Fig 6N), indicating a shift of the Kkv organisation towards “randomness”. Comparing the G-Function SDI value distributions (Fig. 6N’) could not reveal significant differences between the control and the *exp reb* mutants (Kolmogorov-Smirnov D=0.25, *p>0.05*), however this was not surprising as this parameter is less sensitive to change and thus to detect variation amongst samples (Andrey et al., 2010). For this reason, we directly calculated the Nearest Neighbour Distances (NND) of the Kkv puncta from all control and *exp reb* mutant samples and we created the frequency distribution plots for the values obtained (Fig. 6Q). By this approach we were able to detect a significant shift of the distribution frequencies between the two groups (Kolmogorov-Smirnov D=0.2036, *p<0.005*), further supporting the hypothesis of changes in Kkv apical accumulation upon removal of *exp* and *reb*

All these results suggested that *exp/reb* are directly or indirectly required for the proper distribution of Kkv at the apical membrane.

### 7. Exp/Reb and Kkv localise in a complementary pattern

As Exp/Reb proteins localise at the apical membrane (Moussian et al., 2015) and we found that Exp/Reb are required for Kkv apical distribution, we investigated their relative localisations using super-resolution microscopy. Analysis of endogeneous Reb and Kkv proteins in the trachea of wild type embryos indicated that the two proteins in general do not colocalise. Rather, it seemed that they showed a complementary pattern, where Reb accumulated between the Kkv accumulation (Fig 7A). To confirm this observation, we looked at salivary glands expressing Reb. We also found that the Kkv and Reb pattern were complementary and that Reb accumulated particularly where Kkv is low (Fig 7B).

**Figure 7.**
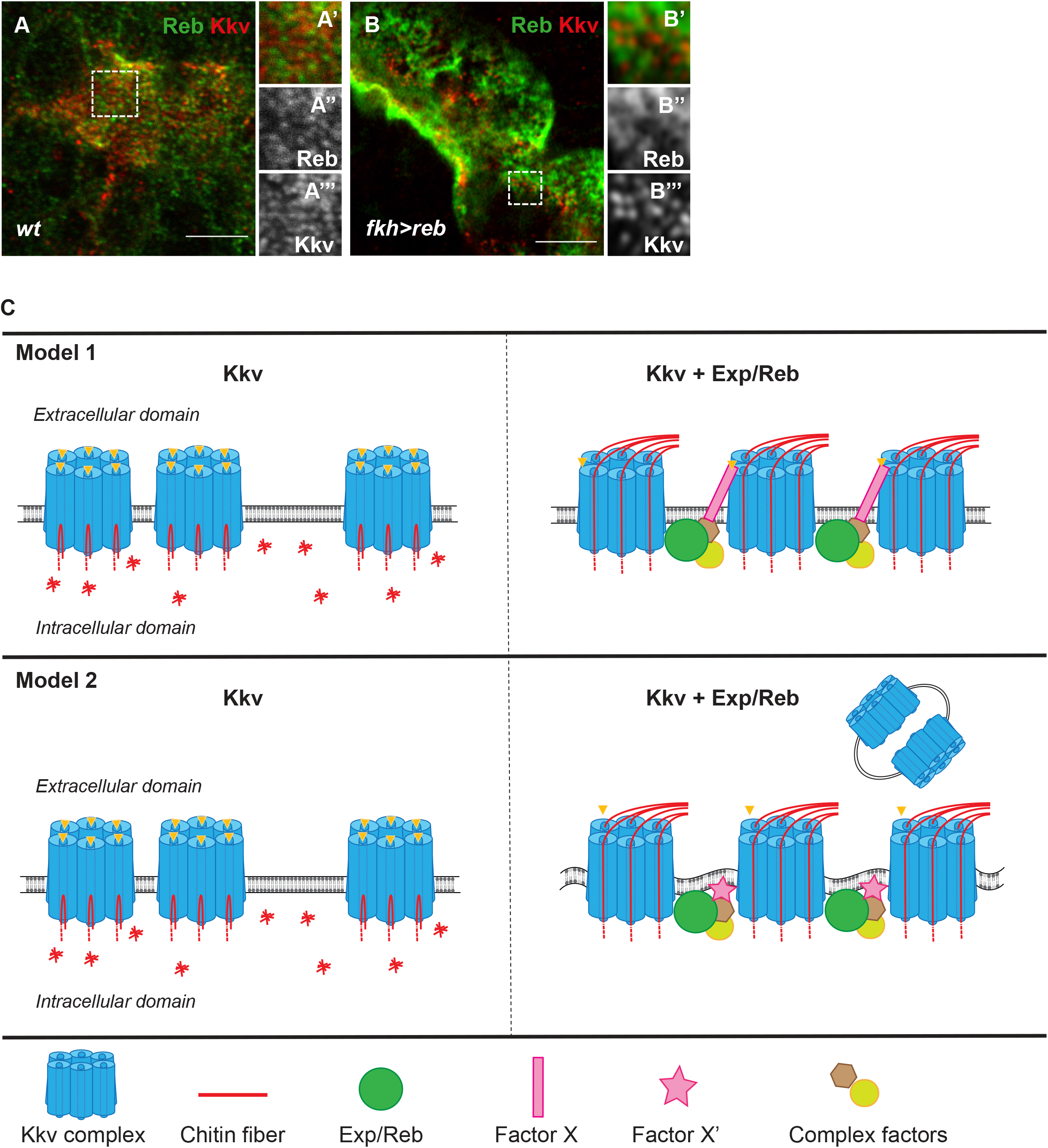
Accumulation of Kkv and Reb. All images are super-resolution single confocal sections. (A) In the trachea of wild type embryos, Reb and Kkv do not colocalise and they show a complementary pattern (A’-A’’’). (B) In salivary gland of embryos expressing Reb, the patterns of Kkv and Reb are complementary. (C) Models for the role of *kkv* and *exp/reb* in chitin deposition. Kkv oligomerises in complexes that localise to the apical membrane (as proposed in (Zhu et al., 2016). In the absence of *exp/reb* activity, Kkv can polymerise chitin from sugar monomers (discontinuous red lines), but it cannot translocate it because the channel is closed. Polymerised chitin remains in the cytoplasm. In addition, Kkv is not homogeneously distributed. Exp/Reb reaches the apical membrane and forms a complex with other proteins. The apical localisation of Exp/Reb complex contributes to Kkv apical distribution. Factors recruited by Exp/Reb (Factor X) can produce a posttranslation or conformational modification to Kkv that promotes translocation of chitin fibers to the extracellular domain (Model1). Alternatively, factors of Exp/Reb complex (Factor X’) can induce changes in membrane composition/curvature that leads to a conformational change in Kkv that opens the channel to translocate chitin. These membrane changes lead to Kkv shedding extracellularly (Model 2). Scale bars 5 μm.

To further investigate whether Kkv and Exp/Reb interact, we performed coimmunoprecipitation experiments. Our αKkv and αReb antibodies recognised Kkv and Reb in embryo extracts (Fig S5). While αKkv antibodies immunoprecipitated the protein with high efficiency (Fig S5A), they did not coimmunoprecipitate Reb (Fig S5B). This negative result may be due to technical difficulties and does not fully rule out an interaction between Kkv and Reb. However, the biochemical and super-resolution analyses do not support a physical interaction between Kkv and Reb.

In summary, our analyses of Exp/Reb and Kkv accumulation suggested that *exp/reb* are indirectly required for the proper distribution of Kkv at the apical membrane.

## DISCUSSION

### Dissection of the roles of conserved motifs in Exp/Reb and Kkv proteins

We generated different transgenic lines under the control of UAS to investigate the function of different conserved domains in Exp/Reb and Kkv proteins. The Gal4/UAS experimental approach allowed us to test whether the mutated proteins are sufficient to promote ectopic or advance chitin deposition (as wild type proteins do), and also whether the mutated proteins can rescue chitin deposition in a mutant background for *exp/reb* or *kkv* respectively (Brand and Perrimon, 1993). However, the Gal4/UAS approach has also different caveats, as it involves the overexpression of the protein, which can bypass or mask the endogenous requirements of specific protein domains, allowing reduced activity mutations to become active.

Our structure-function analysis identified a critical role for the Nα-MH2 domain of Exp/Reb in extracellular chitin translocation. However, the domain was dispensable for chitin polymerisation and for Exp/Reb and Kkv localisation at the apical membrane. The Nα-MH2 domain alone was not able to localise to the membrane and was not functional either. We suggest that Exp/Reb need to localise apically to be active and to promote extracellular chitin translocation. In this context, the Nα-MH2 would be required for interaction/recruitment of other protein/s, presumably in a complex at the apical membrane, which would confer to Kkv the ability to translocate the synthesised polymers. In line with this hypothesis, we identified a new highly conserved motif with a role in Exp/Reb apical accumulation. In our experimental conditions, the CM2 motif was dispensable for *exp/reb* function in extracellular chitin deposition, however, we found that in spite of a clear effect in protein localisation, Exp^ΔCM2^ protein was still able to localise apically, which could account for its activity. We speculate that the massive amount of Exp^ΔCM2^ protein produced in our experimental Gal4/UAS conditions is masking the requirement of the domain for the proper localisation. The generation of CRISPR mutants lacking the CM2 will help to validate this hypothesis in future work.

Kkv is a huge protein with several recognisable domains (Moussian, 2013; Moussian et al., 2005). Our analysis of the conserved WGTRE domain of Kkv indicated that it is required for ER exit. Kkv is produced at the ER and traffics to the membrane where it localises apically to deposit chitin extracellularly. In the case of the yeast chitin synthase Chs3, ER exit requires the activity of the DHHL protein Pfa4 and the dedicated chaperone Chs7 (Lam et al., 2006; Trilla et al., 1999). Hence, ER exit of CHS seems to be a strictly regulated step. In line with this, recent work identified a sarco/endoplasmic reticulum Ca^2+^-ATPase, Serca, that physically interacts with Kkv and regulates chitin deposition (Zhu et al., 2022). The WGTRE motif could directly bind Serca and/or other proteins for ER exit.

The coiled-coil domain of Kkv is predicted to face the extracellular space and may be involved in protein–protein interactions or oligomerisation (Adler, 2019; Merzendorfer, 2006). Our results indicated that Kkv^ΔCC^ behaves as the full length Kkv in several conditions, suggesting that the coiled-coil domain is dispensable for Kkv activity. However, we also found that while the co-expression of *reb* and full length *kkv* in the trachea leads to abnormal tube morphogenesis (due to excessive chitin), the concomitant overexpression of *reb* and *kkv*^*ΔCC*^ resulted in normal tube formation. This is an intriguing behaviour, because the sole overexpression of *reb* in an otherwise wild type condition (i.e. in presence of endogenous *kkv*) produces an abnormal tube morphogenesis. Therefore, the result suggested that *kkv*^*ΔCC*^ is acting to decrease *kkv* endogenous activity. This result would fit with the hypothesis that the CC domain is involved in the oligomerisation of Kkv units. In our model, Kkv^ΔCC^ pure oligomers and Kkv^WT^ oligomers would be fully functional, whereas Kkv^ΔCC^-Kkv^WT^ would not be, maybe due to conformational incompatibilities. In a null mutant background for *kkv*, only Kkv^ΔCC^ oligomers would be present and would support extracellular chitin deposition. In contrast, in a wild type background, Kkv^ΔCC^-Kkv^WT^ inactive oligomers would form, limiting the capacity to promote increased chitin deposition. In support of this hypothesis it has been proposed that CHS may work as oligomers, forming complexes to produce chitin (Gohlke et al., 2017; Maue et al., 2009; Moussian, 2013; Sacristan et al., 2013). On the other hand, we also find that the absence of the CC domain renders a protein that is less efficient to localise apically and that displays a different turnover compared to the wild type Kkv. We propose that this protein would be less active than the wild type form, but again its overexpression would mask this decreased activity. In any case, our results indicate that the CC is involved in protein localisation and trafficking, and point to a correlation between Kkv activity and Kkv trafficking and localisation (see below).

### Kkv and chitin trafficking

Salivary glands proved to be an excellent in vivo test tube to investigate Kkv biology on its own and in relation to chitin deposition. Salivary glands do not deposit chitin in normal conditions. Overexpression of *kkv* (in absence of *exp/reb* activity) promotes polymerisation of chitin that accumulates intracellularly. The simultaneous expression of kkv and exp/reb leads to extracellular chitin deposition in the lumen.

Using a GFP-tagged Kkv protein we followed Kkv trafficking and localisation. We found that Kkv traffics via Golgi, reaches the apical membrane, is then internalised and finally degraded or recycled back to the membrane. A comparable trafficking route has been described for cellulose synthases, CSC (Allen et al., 2021). Also in line with results with CSC, our results indicated a correlation between Kkv activity and Kkv trafficking and localisation (Allen et al., 2021). Kkv protein unable to polymerise chitin did not localize apically and was found diffused in the cytoplasm, strongly suggesting that Kkv polymerisation is important for Kkv localisation/stabilisation at the membrane, as also proposed by other labs (Adler, 2019). On the other hand, fully active Kkv protein able to polymerise and translocate chitin extracellularly becomes strongly enriched apically. We detected some Kkv recycling under these conditions but, strikingly, we also detected the presence of Kkv punctae in the luminal extracellular space. These extracellular Kkv punctae suggested that the protein is shed during or after chitin deposition. Kkv shedding was previously documented in sensory bristles (Adler, 2019; Adler, 2020). Because the GFP tags that allowed us and others (Adler, 2019) to visualise Kkv shedding is cytoplasmic, the results indicate that the whole Kkv protein is shed, rather than cleaved. Membrane proteins can be shed to the extracellular space through exosomes or microvesicles (Tricarico et al., 2017; van Niel et al., 2018). Exosomes derive from endosomal trafficking, but we find that Kkv extracellular vesicles still form when we block endocytosis. Thus, we propose that Kkv is shed through microvesicles when it is actively extruding chitin. Microvesicles arise by outward blebbing and pinching off the plasma membrane, releasing the membrane protein to the extracellular space. Microvesicle formation requires redistribution of membrane lipid and protein components which modulate changes in membrane curvature and rigidity (Tricarico et al., 2017; van Niel et al., 2018). This suggests that conferring ability to Kkv to translocate chitin extracellularly by *exp/reb* activity is linked to membrane reorganisation.

Overexpression of *kkv* in the absence of *exp/reb* activity showed that Kkv is able to polymerise chitin throughout all its trafficking route as we detected exocytic, endocytic and recycling Kkv vesicles containing chitin. This indicates that Kkv is already assembled as a functional synthesising complex in Golgi (but not in ER, as in ER retention conditions no chitin polymerisation is detected), before reaching the membrane, and once it is internalised, as shown for the CSC (Allen et al., 2021). Intriguingly, we also detected the presence of large amounts of membrane-less aggregates of chitin, free of Kkv, in the cytoplasm under these conditions. The exact origin of these chitin aggregates is unclear, however, our observations suggest several mechanisms by which they could form. Chitin could be sorted out from Kkv vesicles by a yet unknown specific and dedicated mechanism. In this case chitin would be polymerised and translocated into the lumen of Kkv vesicles, as proposed for chitosomes (Bartnicki-Garcia, 2006; Bracker et al., 1976; Cohen, 1982). This hypothesis would be consistent with our observation of Kkv vesicles and chitin particles separating. Alternatively, chitin could be polymerised by the catalytic domain of Kkv, which faces the lumen (Adler, 2019), but not translocated and released to the extracellular space unless *exp/reb* activity is present. In this case, non-translocated polymers would be abnormally released intracellularly, remaining in the cytoplasm and organising in aggregates by phase separation (Hyman et al., 2014). Actually, chitin fibrils are insoluble at physiological PH (Elieh-Ali-Komi and Hamblin, 2016). In agreement with this hypothesis, we find a lot of chitin at the subcortical level (where large amounts of Kkv protein localise) and also in close contact with Kkv vesicles. A combination of both mechanisms could also take place.

### Role of Exp/Reb in chitin deposition

It has been proposed that chitin polymerisation occurs in the cytoplasm, where the catalytic domain of CHS localise (Adler, 2019; Merzendorfer, 2006; Merzendorfer, 2011). Our results confirm that chitin is polymerised intracellularly. It has also been proposed that chitin polymerisation is tightly coupled to chitin extrusion (Zhu et al., 2016). In this respect, it has been proposed that the structural organisation of the transmembrane helices of CHS would form a pore or central channel through which the nascent polymer would be translocated (Cohen, 2001; Merzendorfer, 2006; Merzendorfer, 2011; Zhao, 2019; Zhu et al., 2016). Our results, however, indicate that, at least for the case of the chitin synthase Kkv, the polymerisation and extracellular translocation of chitin fibrils are uncoupled, and that while Kkv is sufficient to polymerise the polysaccharide, it cannot translocate it without the activity of *exp/reb*.

What is then the role of *exp/reb* in chitin deposition?

Exp/Reb could potentially play a role in extracellular chitin deposition by binding nascent fibrils polymerised by Kkv and assisting their proper translocation. However, it is unlikely that Exp/Reb could play this role as 1) no chitin binding domain and 2) no transmembrane domain that could form a pore or channel have been identified in these proteins. On the other hand, we do not support either a model in which Exp/Reb form a complex with Kkv, based on the following observations: 1) super-resolution analysis revealed a complementary pattern of Exp/Reb and Kkv at the apical membrane, 2) we could not detect a physical interaction between Reb and Kkv in co-imunoprecipitation experiments, 3) recent work searching for Kkv interacting proteins did not identify Exp or Reb (Duan et al., 2022; Zhu et al., 2022).

Our results indicate that *exp/reb* activity is required for the proper distribution and clustering of Kkv at the apical membrane. It has been proposed that CHS are organised at the plasma membrane in a quaternary structure forming rosettes (Zhu et al., 2016). This hypothesis comes from the comparison between CHS and the closely related CES: both enzymes belong to the β-glycosyltranferase family, produce polymers with similar molecular structure and share several conserved motifs like the catalytic domain QxxRW and several transmembrane domains (Bi et al., 2015; Merzendorfer, 2006). CES organise as hexagonal structures with sixfold symmetries (rosettes). Each rosette is composed of six subunits/lobes, which in turn consist of either six monomeric or three dimeric synthetic units, each capable of polymerising and translocating a single cellulose chain. It has been proposed that differences in the morphology of either the rosettes or their lobes may be responsible for the diversity in cellulose architecture, for instance, dispersed rosettes produce widely spaced cellulose microfibrils, whereas dense regions of complexes synthesize highly aggregated crystalline microfibrils (for reviews see (Allen et al., 2021; Haigler, 2019)). CHS, like Kkv, may function in a similar way (Zhu et al., 2016). In this scenario, we propose that exp/reb controls the distribution/clustering of Kkv rosette-complexes, possibly regulating the spacing by localising in a complementary pattern, which would in turn regulate Kkv activity.

We envision at least two different models, or a combination of them, for a role of *exp/reb* in chitin deposition (Fig 7C). In both models Kkv has a closed conformation in the absence of Exp/Reb that prevents chitin translocation, and Exp/Reb regulate the apical distribution of Kkv. In our first model, Exp/Reb are required to organise a complex (through its Nα-MH2 domain) that reaches the apical membrane (through its CM2). One of the components of the complex, Factor X, interacts directly with Kkv promoting a posttranscriptional modification or a conformational change in Kkv that activates its translocation activity. In the absence of Exp/Reb, Factor X would not reach the membrane and Kkv would remain plugged or clogged preventing the translocation of the chitin fibrils. In a second model, Exp/Reb are required (in collaboration with its complex) to promote a favourable membrane environment (e.g. curvature, membrane composition) for the proper integration/insertion/arrangement of Kkv rosette complexes. This arrangement of the complex is required to impose a particular conformational organisation in the rosette that opens the pore of the Kkv units, which otherwise are plugged or clogged. In agreement with this second model, it has been suggested that the Kkv interacting protein Ctl2 could regulate membrane phospholipid composition and increase Kkv activity (Duan et al., 2022). In addition, this model would fit with the putative role of membrane curvature and composition and Kkv shedding as microvesicles (Tricarico et al., 2017; van Niel et al., 2018). A similar mechanism has been shown for the mechanosensor Piezo, where the conducting conformation of its pore (open or closed) can be controlled by local changes in membrane curvature and tension (Jiang et al., 2021; Yang et al., 2022).

In summary, our results point to an absolut requirement for *exp/reb* in regulating the capacity of Kkv to translocate the nascent chitin fibrils, likely regulating its distribution and conformational organisation at the apical membrane. This reveals the existence of an extrinsic mechanism of regulation of CHS which seems to be conserved during evolution, as orthologs for Exp have been identified in arthropods and nematodes.

## ACKNOWLEDGEMENTS

We thank E. Rebollo from the IBMB-PCB Molecular Imaging Platform for imaging and N. Giakoumakis and S. Tosi from IRB-Advanced Digital Microscopy for technical help and discussions on quantifications. We thank N. Martín for technical help and generation of Kkv antibody. We acknowledge the Bloomington Stock Centre and the Developmental Studies Hybridoma Bank for fly lines and antibodies. We thank T. Tanaka, S. Hayashi and D. Andrew for kindly providing flies and antibodies. Thanks also go to the members of the Llimargas and Casanova labs for helpful discussions, and to J. Casanova and M. Furriols for critical reading of the manuscript.

EDG was funded by a FPI Fellowship of the Spanish Ministerio de Ciencia e Innovación (BES-2016-076723). This work was funded by the Spanish Ministerio de Ciencia e Innovación (MICINN, https://www.ciencia.gob.es/, grants numbers BFU-2015-68098-P and PGC2018-098449-B-I00 to ML). The funders had no role in study design, data collection and analysis, decision to publish, or preparation of the manuscript. The authors declare no competing interests

## Author contributions

Conceptualization: EDG, ML Methodology: EDG, PG Formal analysis: EDG, PG, ML

Investigation: EDG, PG, MLE, ML Writing – original draft: ML

Writing –review & editing: EDG, PG, MLE, ML Supervision: ML

Funding acquisition: ML

## MATERIALS AND METHODS

### Resources Table

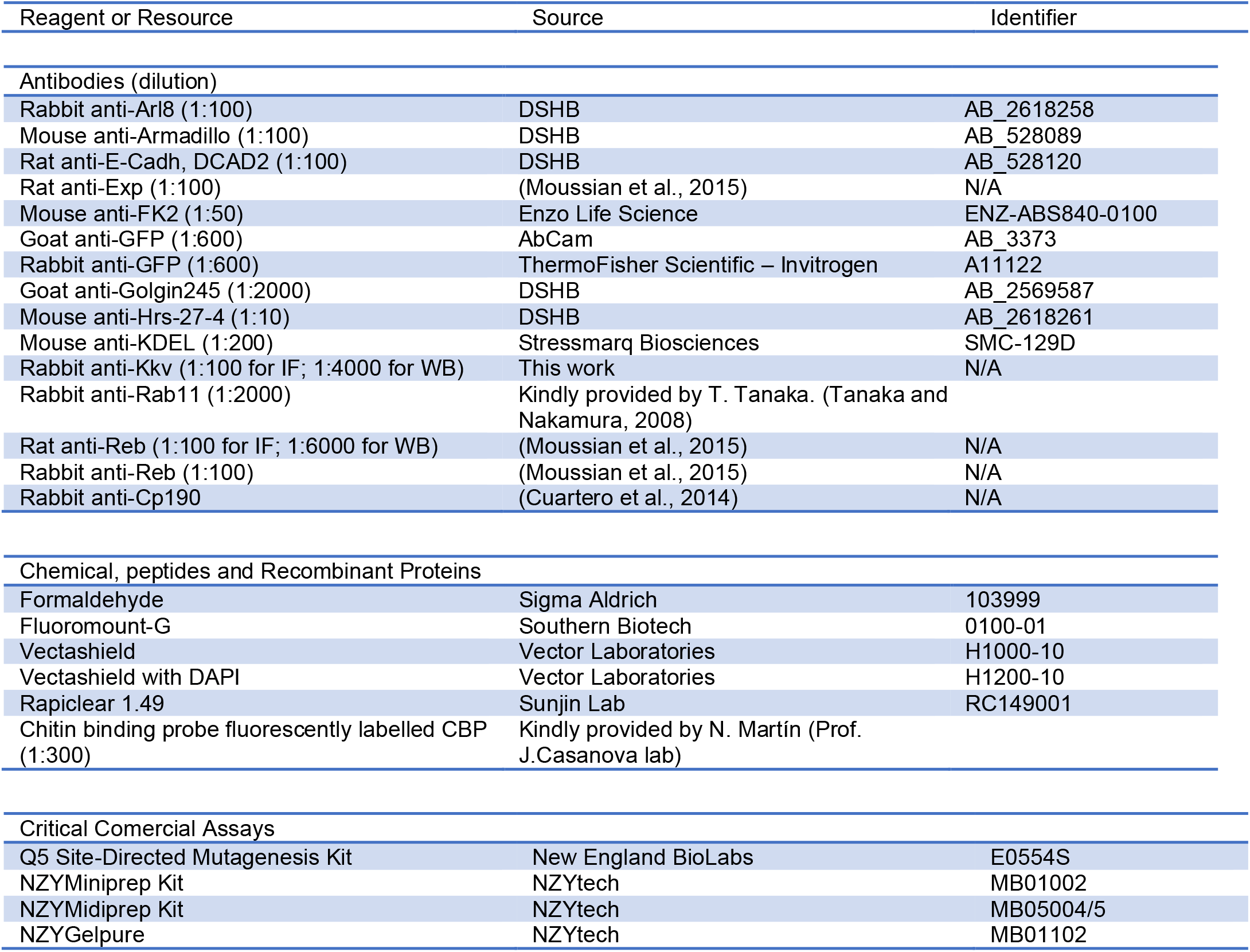
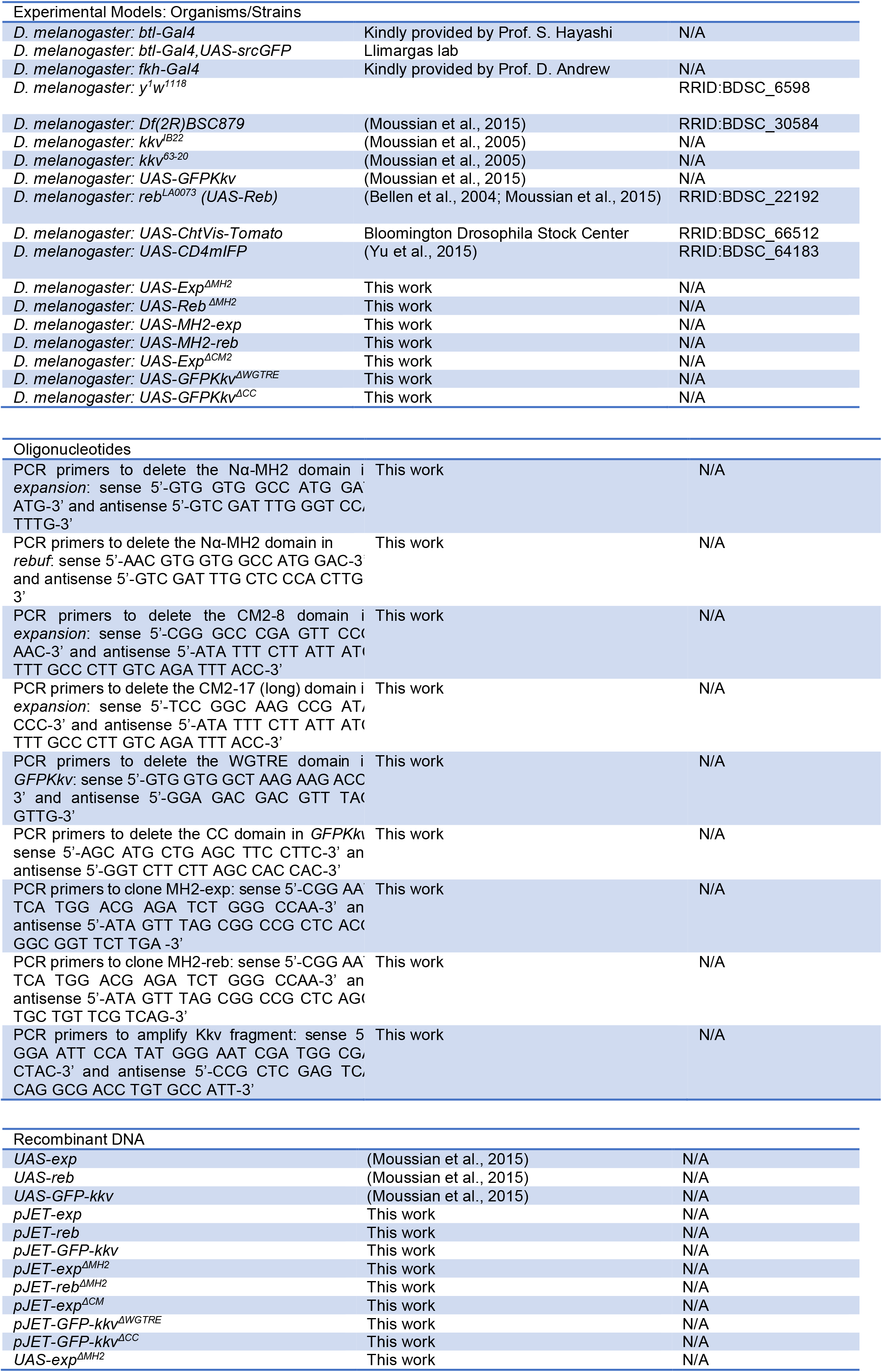
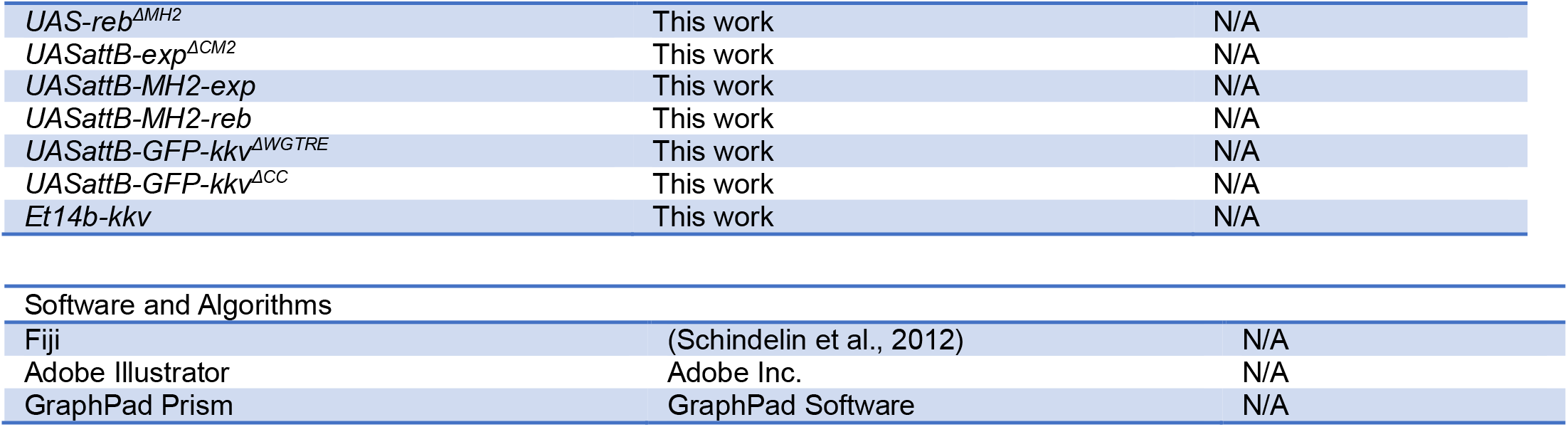

### *Drosophila* strains and Maintenance

All *Drosophila* strains were raised at 25ºC under standard conditions. The strain y^1^w^1118^ was used as wild type (wt). Balancer chromosomes were used to follow the mutations and constructs of interest in the different chromosomes. For overexpression experiments, we used the Gal4 drivers *btlGal4* (in all tracheal cells) and *fkhGal4* (in salivary glands). The overexpression and rescue experiments were performed using the Gal4/UAS system (Brand and Perrimon, 1993). To maximise the expression of the transgenes, crosses were kept at 29°C. The fly strains used are listed in the “Resource table: Experimental models”.

### Immunohistochemistry

Embryos were stained following standard protocols. Embryos were staged as described (Campos-Ortega and Hartenstein, 1985). Embryos were fixed in 4% formaldehyde (Sigma-Aldrich) in PBS1x-Heptane (1:1) for 10 min for Ecad staining and for 20 min for the rest. Embryos transferred to new tubes were washed in PBT-BSA blocking solution and shaken in a rotator device at room temperature. Embryos were incubated with the primary antibodies in PBT-BSA overnight at 4ºC. Secondary antibodies diluted in PBT-BSA (and for the CBP staining) were added after washing and were incubated at room temperature for 2–5 h in the dark. Embryos were washed, mounted on microscope glass slides and covered with thin glass slides. The primary antibodies used are listed in the “Key Resource table: Antibodies”. Cy3-, Cy2-and Cy5-conjugated secondary antibodies (Jackson ImmunoResearch) were used at 1:300.

### Image acquisition

Fluorescence confocal images of fixed embryos were obtained with Leica TCS-SPE system using 20x and 63x (1,40-0,60 oil) objectives (Leica). For super-resolution images two different systems were used: Elyra PS1-Airyscan (Zeiss) from the IRB-Advanced Digital Microscopy Core Facilty and Drangofly 505 (Andor) from IBMB-Molecular Imaging Platform; in both cases a 100x (1,40-0,60 oil) objective was used. The latter system was used also to perform life imaging. In this case, dechorionated embryos were mounted and lined up on a Menzel-Glaser coverslip with oil 10-S Voltalef (VWR) and covered with a membrane (YSI membrane kit). In all movies we used 63x (1,40-0,60 oil) objective. To visualise time-lapse movies, single sections were used. Fiji (ImageJ) was used for measurement and adjustment. Otherwise indicated in the text, confocal images are maximum-intensity projections of Z-stack sections. Figures were assembled with Adobe Illustrator.

### Generation of UAS constructs

For the generation of new recombinant DNA we used pUAST-exp, -reb and -GFPkkv (Moussian et al., 2015). We digested the DNA from the vector using the following couples of restriction enzymes (New England BioLabs, NEB): EcoRI/XhoI for *exp*, EcoRI/NotI for *reb* and XhoI/XbaI for *GFP-kkv* and we cloned them in the vector pJET1.2.

The constructs UAS-exp^ΔMH2^, -reb^ΔMH2^, -exp^ΔCM2^, -GFP-kkv^ΔWGTRE^ and -GFP-kkv^ΔCC^ were obtained by directed deletion. To delete specific regions from the DNA, “Q5 Site-Directed Mutagenesis Kit” (NEB) was used. The kit comprehended material to perform PCR, ligation of the fragments and transformation of NEB-α competent cells. Deletions are created by designing primers that flank the region to be deleted, then we performed a PCR obtaining a linear double filament of DNA composed by the original pJET1.2 vector and the flanking regions of the deleted sequence. Upon ligation of the fragment, competent cells were transformed and plated in selective plates. Miniprep to obtain DNA were performed using the kit NZYtech and the DNA was sequenced through the platform Eurofins Genomics. Finally, the new mutated DNA was digest using the restriction enzymes described above and cloned in pUAST vector for exp^ΔMH2^ and reb^ΔMH2^ and in pUAST-attB vector for all the other DNAs. After performing a midiprep (NZYtech), the DNAs were injected in embryos y^1^w^1118^ by the “Drosophila injection Service” of the “Institute for Research in Biomedicine” (IRB, Barcelona) and by the “Transgenesis Service” of the “Centro de Biología Molecular Severo Ochoa” (CBM, Madrid). The primers used are listed in the “Resource table: Oligonucleotides”. UAS-MH2-exp and UAS-MH2-reb were obtained amplifying the MH2 region respectively from pJET-exp and pJET-reb and cloning the fragment in the pUAST-attB vector using in both cases EcoRI/NotI. The primers used are listed in the “Key Resource table: Oligonucleotides”.

### Generation of antibodies

To generate polyclonal antibody against Kkv, fragments were amplified by PCR using the primers listed in “Resource table: Oligonucleotides”. We used the restriction sites NdeI/XhoI. The amplified fragments were cloned into the expression vector pET14b (Novagen). The resulting positive clones were used to transform BL2 (C41) cells (Novagen) for protein expression. Cells were induced with 1 mM IPTG and proteins were expressed at 37ºC during 2 hours. The positive clones were selected and the recombinant proteins (22 KDa) fused with a His tag were purified through a column of nickel (Quiagen) in denaturalising conditions (8 M urea). The purified proteins were used to inject rabbits by the facility CID-CSIC-Production of antibodies (Barcelona).

### Quantification and statistical analysis

Data from quantifications was imported and treated in the Excel software and/or in GraphPad Prism 9.0.0, where graphics were finally generated. Graphics shown are scatter dot plots, where bars indicate the mean and the standard deviation. Statistical analyses comparing the mean of two groups of quantitative continuous data were performed in GraphPad Prism 9.0.0 using unpaired two-tailed student’s t-test applying Welch’s correction. Differences were considered significant when p < 0.05. Significant differences are shown in the graphics as: *p < 0.05, **p < 0.01, ***p < 0.001, ****p < 0.0001. n.s. means not statistically significant. Sample size (n) is provided in the figures or legends. The non-parametric Kolmogorov-Smirnov test was used for the comparisons of frequency distributions.

### Image analyses

#### Apical/Basal localisation

On single section images, we measured the Integrated Density (IntDen) of the apical and basal region using the free hand line tool and the measure tool of Fiji software. Each value of the graphs represents the ratio between the IntDen of the apical region and the IntDen of its respective basal region.

#### Number of vesicles/particles

Max Intensity projections of 13 sections (0,29 µm each) and Fiji software were used. The subtract background tool, the threshold tool and the watershed tool were applied to create binarised masks of the vesicles. Numbers of vesicles were counted using the Analyse Particles tool and the parameters were set to 0,02-1,1 µm^2^ size, 0-1 circularity; a mask of the result was obtained.

#### Colocalisation of vesicles/particles

The “And” operation of the Image Calculator tool of Fiji software was applied between masks obtained as a result of the “Number of vesicles” process. The resulting image was analysed through the Analyse Particle tool as described above.

#### Analysis of Kkv distribution

For the Kkv distribution analysis, we used maximum intensity projections of the same number of stacks for all cells, to create binarized masks of the apical area defined by the Arm marker. For the detection of the Kkv puncta and the subsequent creation of a second binary image we used the Find Maxima function of the Fiji software, with the output set to Maxima with Tolerance. The two binary images created were used as an input for the Spatial Statistics 2D/3D Fiji plugin (Andrey et al., 2010; Ollion et al., 2013) and the parameters were set to: 10000 number of points for the F-Function, 200 random point pattern generation for the average CDF and SDI, hardcore distance of 0.08 to 0.18 and confidence limit for the CDF at 5%. For the calculation of the NND, the binary images of the Kkv spots created previously, were used as an input for the Nearest Neighbor Distances Calculation with ImageJ plugin of the Fiji software. For the analysis of the results obtained, the Kolmogorov-Smirnov test was used to assess the sample distribution within the populations.

### Co-immunoprecipitation assay

Assays were performed with extracts prepared from *Drosophila* embryos that were lysed in RIPA buffer (50 mM Tris-HCl pH8,150 mM NaCl, 0.1% SDS, 0.5% sodium deoxycholate,1% Triton X-100, 1mM PMSF and protease inhibitors (cOmplete Tablets, Roche). Extracts were immunoprecipitated using anti-Kkv antibodies or a mock antibody (anti-CP190), followed by incubation with Protein G Dynabeads (Invitrogen). Immunoprecipitates were washed with RIPA buffer and analysed by Western blot using anti-Kkv or anti-Reb antibodies and the Immobilon ECL reagent (Millipore).

## FIGURE LEGENDS

**Figure S1.**
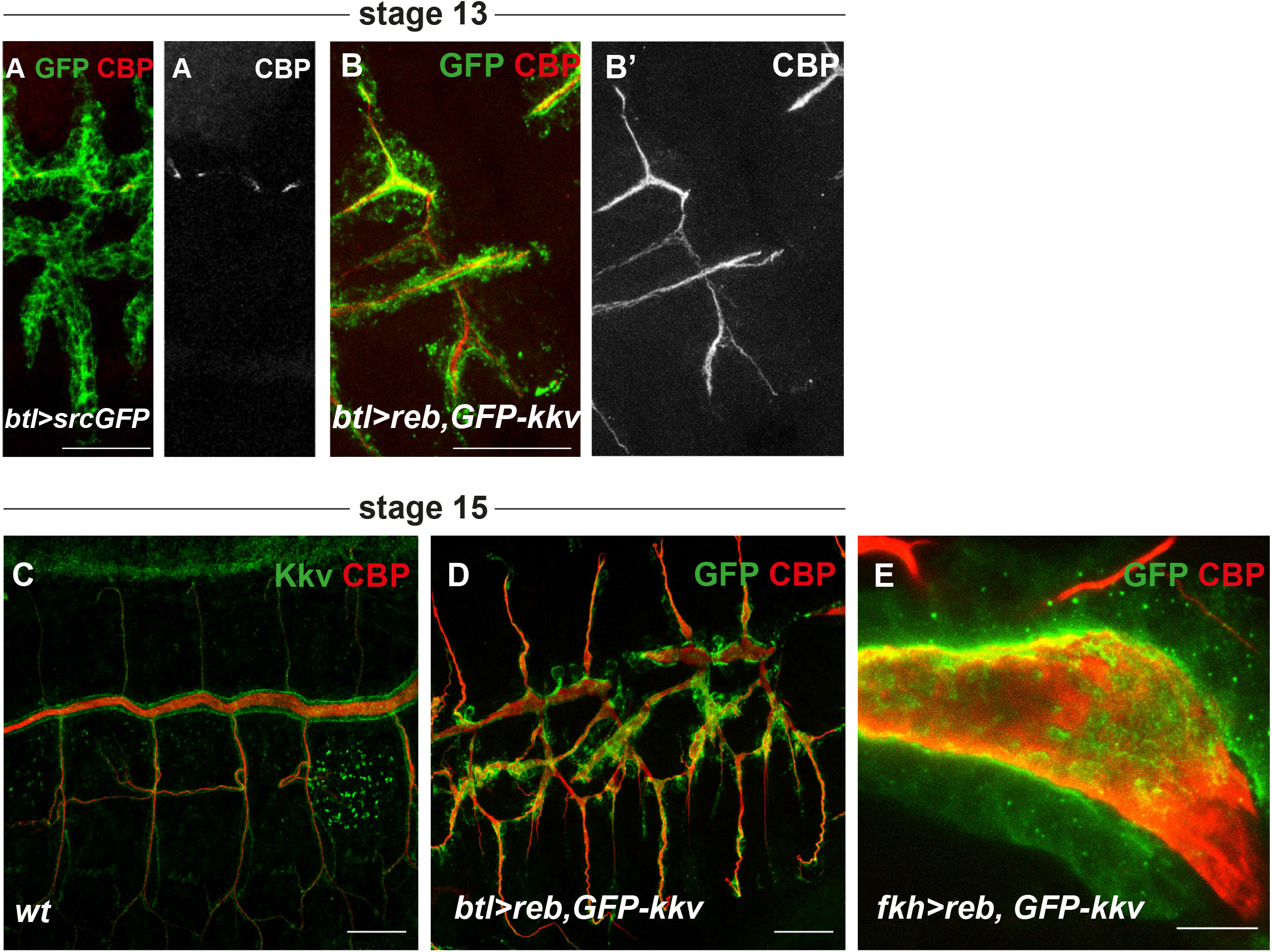
Effects of the co-expression of Reb and GFP-Kkv expression. All images correspond to projections of confocal sections. (A-D) In trachea, the simultaneous overexpression of *reb* and *GFP-kkv* anticipates chitin deposition (compare A and B) and produces morphogenetic defects (compare C and D). (E) In salivary glands, the co-expression of *reb* and *GFP-kkv* promotes accumulation of chitin in the lumen. Scale bars A-D 25 μm, E 10 μm.

**Figure S2.**
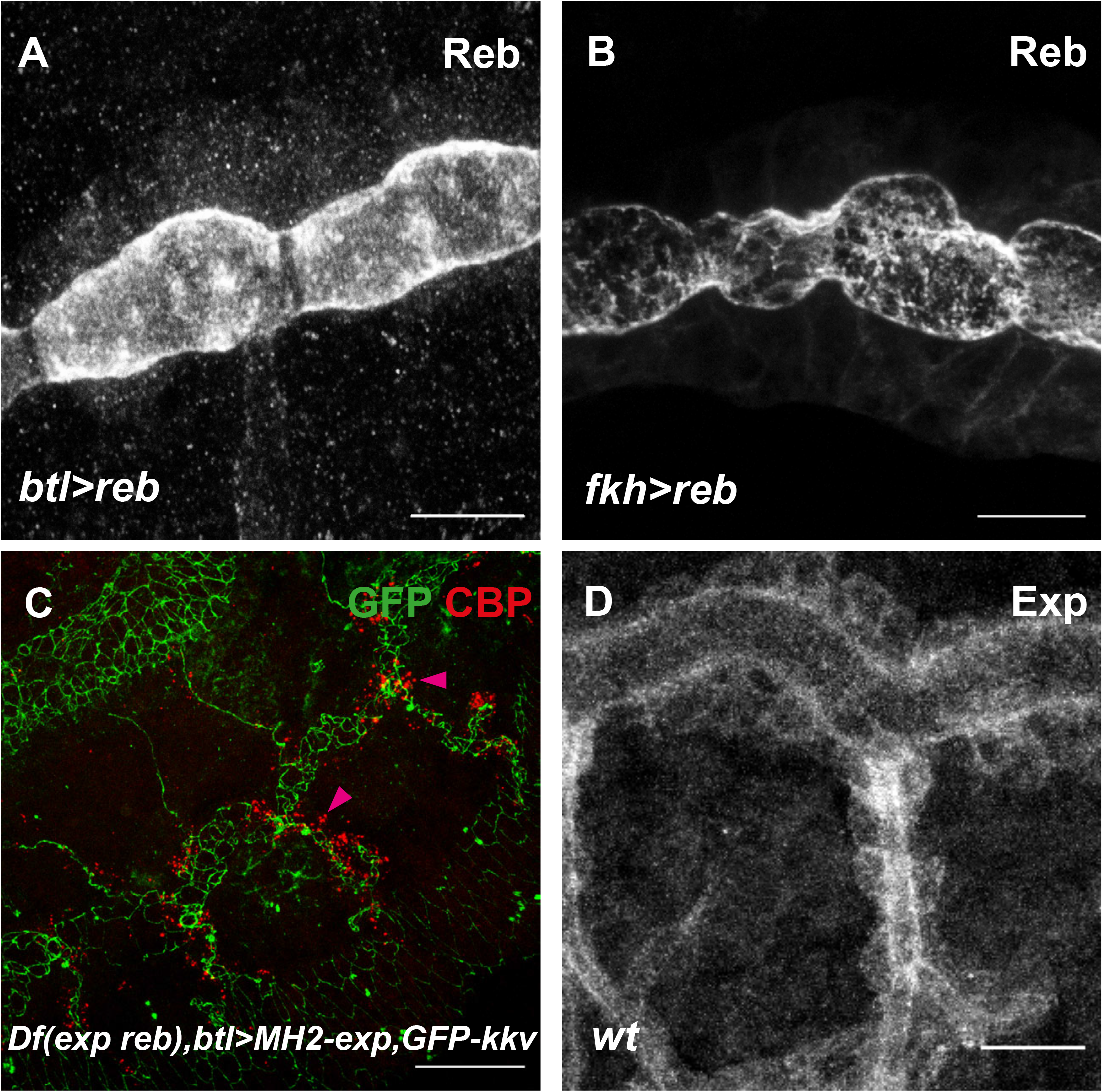
Effects of the expression of Exp/Reb and GFP-Kkv. All images are projections of confocal sections. (A,B) Overexpressed full-length Reb localises mainly apically in trachea (A) and in salivary glands (B). (C) The simultaneous overexpression of *MH2-exp* and *GFP-kkv* does not rescue the absence of extracellular chitin deposition and it produces intracellular chitin vesicles (pink arrowheads). (D) Endogenous Exp localises mainly apically in trachea, although a bit of the protein can be detected intracellularly. Scale bars 10 μm.

**Figure S3.**
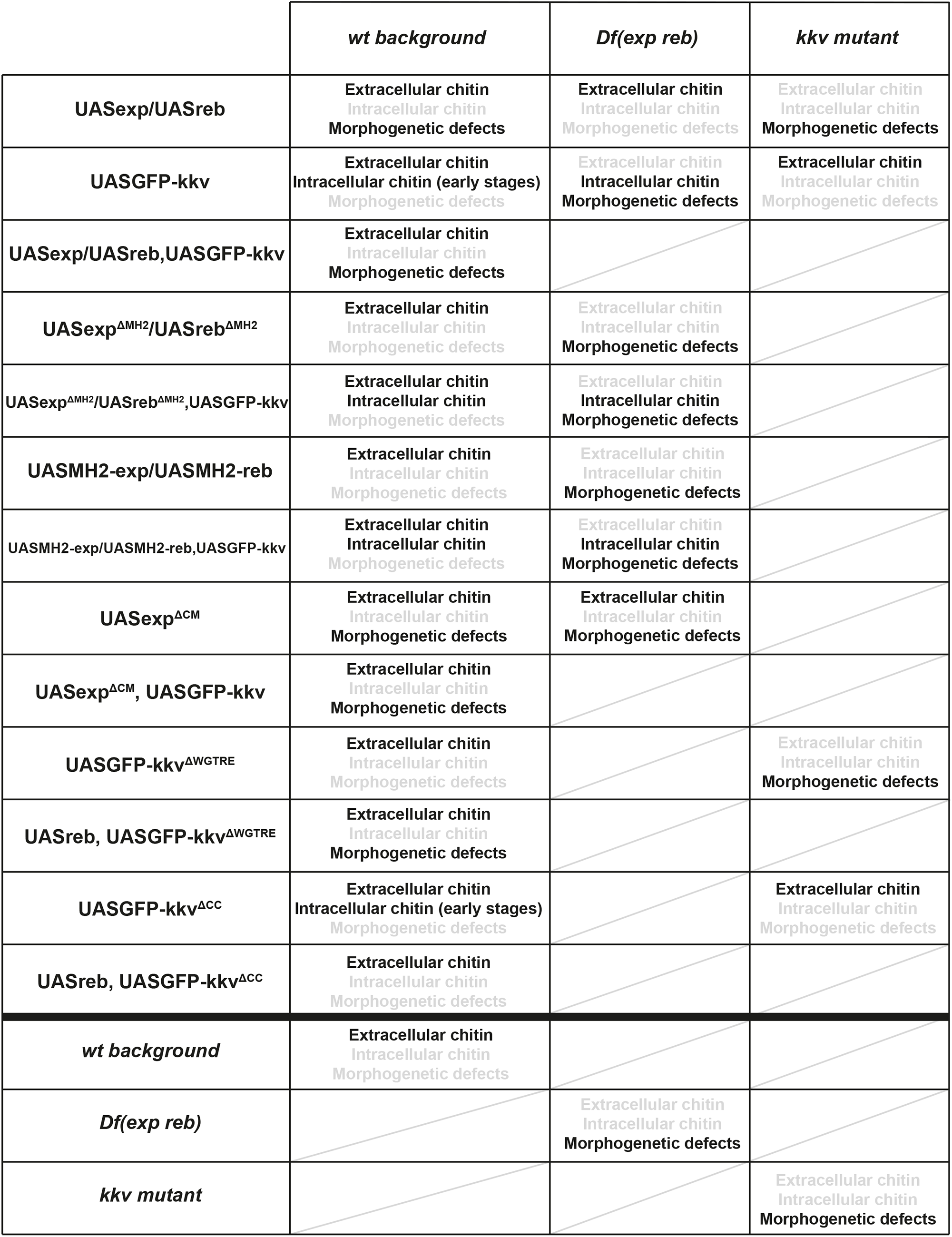
Summary of described phenotypes. Summary of phenotypes of the different UAS constructs in wild type (wt), *exp reb* mutant and *kkv* mutant embryos in different overexpression conditions. The phenotypes observed are indicated in black, and light grey indicates absence of the phenotype

**Figure S4.**
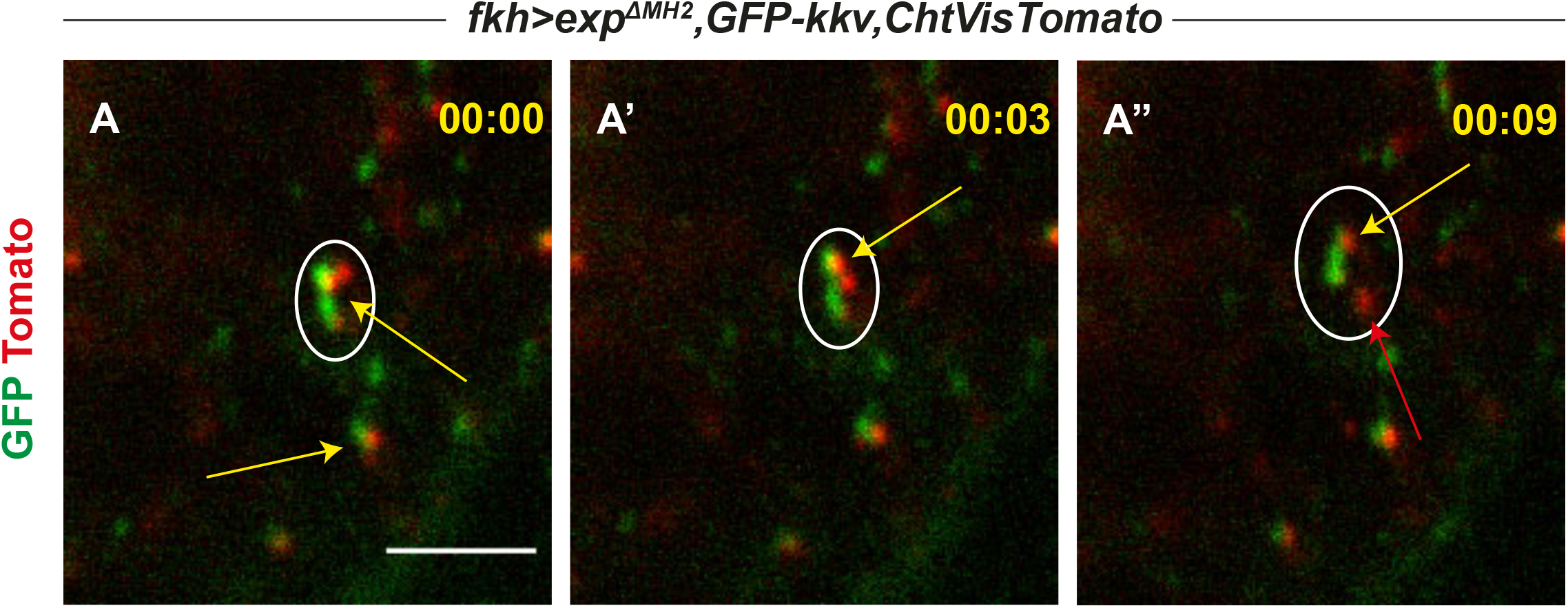
Frames from live imaging movie 2 show that common GFP-Kkv and chitin punctae (yellow arrow) can separate from each other; many GFP-Kkv (green arrow) and chitin puncta (red arrow) do not colocalise. Scale bar 5 μm.

**Figure S5.**
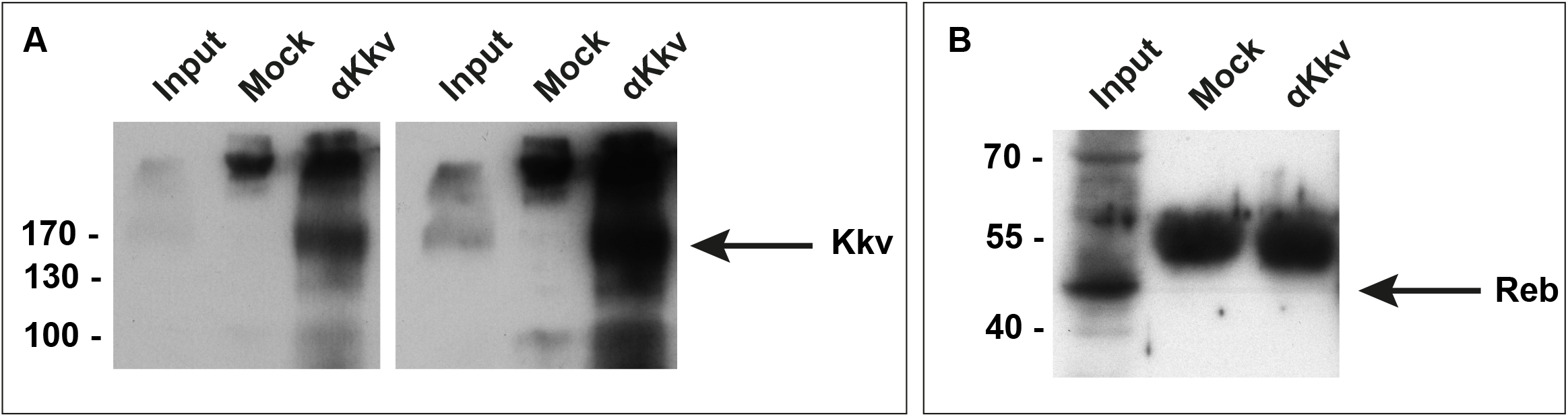
Co-IP. Western blot using αKkv (A, two different exposure times are shown) or αReb (B) of embryo extracts that were subjected to immunoprecipitation with αKkv or an unrelated antibody (mock). Input correspond to 7.5 % of the immunoprecipitated material. The position of MW markers (in kDa) is indicated.

**Movies 1 and 2. GFP-kkv vesicles and chitin punctae**.

Salivary glands of stage 15 embryos carrying *GFP-kkv, ChtVisTomato* and *exp*^*ΔMH2*^ visualised from a lateral view using Drangofly 505 (Andor) with 63x oil objective and a 2x zoom. Images were taken every 3 seconds in one single Z-stack during two minutes for movie 1 and during 1 minute and 30 seconds for movie 2. The movies show a chitin particle detaching from a GFP-kkv vesicle.

## REFERENCES

Adler, P.N. 2019. The localization of chitin synthase mediates the patterned deposition of chitin in developing Drosophila bristles. BioRxiv.

Adler, P.N. 2020. Short distance non-autonomy and intercellular transfer of chitin 1 synthase in Drosophila. BioRxiv.

Allen, H., D. Wei, Y. Gu, and S. Li. 2021. A historical perspective on the regulation of cellulose biosynthesis. Carbohydrate polymers. 252:117022.

Andrey, P., K. Kieu, C. Kress, G. Lehmann, L. Tirichine, Z. Liu, E. Biot, P.G. Adenot, C. Hue-Beauvais, N. Houba-Herin, V. Duranthon, E. Devinoy, N. Beaujean, V. Gaudin, Y. Maurin, and P. Debey. 2010. Statistical analysis of 3D images detects regular spatial distributions of centromeres and chromocenters in animal and plant nuclei. PLoS computational biology. 6:e1000853.

Arpon, J., K. Sakai, V. Gaudin, and P. Andrey. 2021. Spatial modeling of biological patterns shows multiscale organization of Arabidopsis thaliana heterochromatin. Scientific reports. 11:323.

Bartnicki-Garcia, S. 2006. Chitosomes: past, present and future. FEMS yeast research. 6:957–965.

Beich-Frandsen, M., E. Aragon, M. Llimargas, J. Benach, A. Riera, J. Pous, and M.J. Macias. 2015. Structure of the N-terminal domain of the protein Expansion: an ‘Expansion’ to the Smad MH2 fold. Acta Crystallogr D Biol Crystallogr. 71:844–853.

Bellen, H.J., R.W. Levis, G. Liao, Y. He, J.W. Carlson, G. Tsang, M. Evans-Holm, P.R. Hiesinger, K.L. Schulze, G.M. Rubin, R.A. Hoskins, and A.C. Spradling. 2004. The BDGP gene disruption project: single transposon insertions associated with 40% of Drosophila genes. Genetics. 167:761–781.

Bi, Y., C. Hubbard, P. Purushotham, and J. Zimmer. 2015. Insights into the structure and function of membrane-integrated processive glycosyltransferases. Current opinion in structural biology. 34:78–86.

Bracker, C.E., J. Ruiz-Herrera, and S. Bartnicki-Garcia. 1976. Structure and transformation of chitin synthetase particles (chitosomes) during microfibril synthesis in vitro. Proc Natl Acad Sci U S A. 73:4570–4574.

Brand, A.H., and N. Perrimon. 1993. Targeted gene expression as a means of altering cell fates and generating dominant phenotypes. Development. 118:401–415.

Campos-Ortega, A.J., and V. Hartenstein. 1985. The Embryonic Development of Drosophila Melanogaster. Springer-Verlag. New York:10–84.

Casadidio, C., D.V. Peregrina, M.R. Gigliobianco, S. Deng, R. Censi, and P. Di Martino. 2019. Chitin and Chitosans: Characteristics, Eco-Friendly Processes, and Applications in Cosmetic Science. Marine drugs. 17.

Cohen, E. 1982. In vitro chitin synthesis in an insect: formation and structure of microfibrils. Eur J Cell Biol. 26:289–294.

Cohen, E. 2001. Chitin synthesis and inhibition: a revisit. Pest Manag Sci. 57:946–950.

Cuartero, S., U. Fresan, O. Reina, E. Planet, and M.L. Espinas. 2014. Ibf1 and Ibf2 are novel CP190-interacting proteins required for insulator function. EMBO J. 33:637–647.

Duan, Y., W. Zhu, X. Zhao, H. Merzendorfer, J. Chen, X. Zou, and Q. Yang. 2022. Choline transporter-like protein 2 interacts with chitin synthase 1 and is involved in insect cuticle development. Insect Biochem Mol Biol. 141:103718.

Elieh-Ali-Komi, D., and M.R. Hamblin. 2016. Chitin and Chitosan: Production and Application of Versatile Biomedical Nanomaterials. International journal of advanced research. 4:411–427.

Fujimuro, M., H. Sawada, and H. Yokosawa. 1994. Production and characterization of monoclonal antibodies specific to multi-ubiquitin chains of polyubiquitinated proteins. FEBS Lett. 349:173–180.

Gohlke, S., S. Muthukrishnan, and H. Merzendorfer. 2017. In Vitro and In Vivo Studies on the Structural Organization of Chs3 from Saccharomyces cerevisiae. International journal of molecular sciences. 18.

Haigler, C.H., Roberts, A.W. 2019. Structure/function relationships in the rosette cellulose synthesis complex illuminated by an evolutionary perspective. Cellulose. 26:277–247.

Hyman, A.A., C.A. Weber, and F. Julicher. 2014. Liquid-liquid phase separation in biology. Annu Rev Cell Dev Biol. 30:39–58.

Jiang, Y., X. Yang, J. Jiang, and B. Xiao. 2021. Structural Designs and Mechanogating Mechanisms of the Mechanosensitive Piezo Channels. Trends in biochemical sciences. 46:472–488.

Lam, K.K., M. Davey, B. Sun, A.F. Roth, N.G. Davis, and E. Conibear. 2006. Palmitoylation by the DHHC protein Pfa4 regulates the ER exit of Chs3. J Cell Biol. 174:19–25.

Liu, X., A.M.W. Cooper, J. Zhang, and K.Y. Zhu. 2019. Biosynthesis, modifications and degradation of chitin in the formation and turnover of peritrophic matrix in insects. J Insect Physiol. 114:109–115.

Maue, L., D. Meissner, and H. Merzendorfer. 2009. Purification of an active, oligomeric chitin synthase complex from the midgut of the tobacco hornworm. Insect Biochem Mol Biol. 39:654–659.

Merzendorfer, H. 2006. Insect chitin synthases: a review. J Comp Physiol B. 176:1–15.

Merzendorfer, H. 2011. The cellular basis of chitin synthesis in fungi and insects: common principles and differences. Eur J Cell Biol. 90:759–769.

Merzendorfer, H., and L. Zimoch. 2003. Chitin metabolism in insects: structure, function and regulation of chitin synthases and chitinases. J Exp Biol. 206:4393–4412.

Moussian, B. 2013. The apical plasma membrane of chitin-synthesizing epithelia. Insect Sci. 20:139–146.

Moussian, B., A. Letizia, G. Martinez-Corrales, B. Rotstein, A. Casali, and M. Llimargas. 2015. Deciphering the genetic programme triggering timely and spatially-regulated chitin deposition. PLoS Genet. 11:e1004939.

Moussian, B., H. Schwarz, S. Bartoszewski, and C. Nusslein-Volhard. 2005. Involvement of chitin in exoskeleton morphogenesis in Drosophila melanogaster. J Morphol. 264:117–130.

Ollion, J., J. Cochennec, F. Loll, C. Escude, and T. Boudier. 2013. TANGO: a generic tool for high-throughput 3D image analysis for studying nuclear organization. Bioinformatics. 29:1840–1841.

Ostrowski, S., H.A. Dierick, and A. Bejsovec. 2002. Genetic control of cuticle formation during embryonic development of Drosophila melanogaster. Genetics. 161:171–182.

Sacristan, C., J. Manzano-Lopez, A. Reyes, A. Spang, M. Muniz, and C. Roncero. 2013. Oligomerization of the chitin synthase Chs3 is monitored at the Golgi and affects its endocytic recycling. Molecular microbiology. 90:252–266.

Schindelin, J., I. Arganda-Carreras, E. Frise, V. Kaynig, M. Longair, T. Pietzsch, S. Preibisch, C. Rueden, S. Saalfeld, B. Schmid, J.Y. Tinevez, D.J. White, V. Hartenstein, K. Eliceiri, P. Tomancak, and A. Cardona. 2012. Fiji: an open-source platform for biological-image analysis. Nat Methods. 9:676–682.

Tanaka, T., and A. Nakamura. 2008. The endocytic pathway acts downstream of Oskar in Drosophila germ plasm assembly. Development. 135:1107–1117.

Tricarico, C., J. Clancy, and C. D’Souza-Schorey. 2017. Biology and biogenesis of shed microvesicles. Small GTPases. 8:220–232.

Trilla, J.A., A. Duran, and C. Roncero. 1999. Chs7p, a new protein involved in the control of protein export from the endoplasmic reticulum that is specifically engaged in the regulation of chitin synthesis in Saccharomyces cerevisiae. J Cell Biol. 145:1153–1163.

Tsai, Y.C., and A.M. Weissman. 2010. The Unfolded Protein Response, Degradation from Endoplasmic Reticulum and Cancer. Genes & cancer. 1:764–778.

van Niel, G., G. D’Angelo, and G. Raposo. 2018. Shedding light on the cell biology of extracellular vesicles. Nat Rev Mol Cell Biol. 19:213–228.

Yang, X., C. Lin, X. Chen, S. Li, X. Li, and B. Xiao. 2022. Structure deformation and curvature sensing of PIEZO1 in lipid membranes. Nature. 604:377–383.

Younes, I., and M. Rinaudo. 2015. Chitin and chitosan preparation from marine sources. Structure, properties and applications. Marine drugs. 13:1133–1174.

Yu, D., M.A. Baird, J.R. Allen, E.S. Howe, M.P. Klassen, A. Reade, K. Makhijani, Y. Song, S. Liu, Z. Murthy, S.Q. Zhang, O.D. Weiner, T.B. Kornberg, Y.N. Jan, M.W. Davidson, and X. Shu. 2015. A naturally monomeric infrared fluorescent protein for protein labeling in vivo. Nat Methods. 12:763–765.

Zhao, X., Zhang, J., Zhu, K.Y. 2019. Chito-Protein Matrices in Arthropod Exoskeletons and Peritrophic Matrices. In Extracellular Sugar-Based Biopolymers Matrices. Biologically-Inspired Systems. Vol. 12. E. Cohen, Merzendorfer, H. (eds), editor. Springer, Cham.

Zhu, K.Y., H. Merzendorfer, W. Zhang, J. Zhang, and S. Muthukrishnan. 2016. Biosynthesis, Turnover, and Functions of Chitin in Insects. Annu Rev Entomol. 61:177–196.

Zhu, W., Y. Duan, J. Chen, H. Merzendorfer, X. Zou, and Q. Yang. 2022. SERCA interacts with chitin synthase and participates in cuticular chitin biogenesis in Drosophila. Insect Biochem Mol Biol. 145:103783.

